# Quantifying and modelling the acquisition and retention of lumpy skin disease virus by haematophagus insects reveals clinically but not subclinically-affected cattle are promoters of viral transmission and key targets for control of disease outbreaks

**DOI:** 10.1101/2020.06.18.154252

**Authors:** Beatriz Sanz-Bernardo, Ismar R. Haga, Najith Wijesiriwardana, Sanjay Basu, Will Larner, Adriana V. Diaz, Zoë Langlands, Eric Denison, Joanne Stoner, Mia White, Christopher Sanders, Philippa C. Hawes, Anthony J. Wilson, John Atkinson, Carrie Batten, Luke Alphey, Karin E. Darpel, Simon Gubbins, Philippa M. Beard

**Affiliations:** The Pirbright Institute, Ash Road, Pirbright, Surrey, UK; The Roslin Institute, Easter Bush, Edinburgh, UK; MSD Animal Health, Walton Manor, Walton, Milton Keynes, UK

**Keywords:** poxvirus, lumpy skin disease, transmission, mosquitoes, flies, midges, basic reproduction number, vector, control, *Aedes aegypti*, *Culex quinquefasciatus*, *Stomoxys calcitrans*, *Culicoides nubeculosus*

## Abstract

Lumpy skin disease virus (LSDV), a poxvirus that causes severe disease in cattle, has in the last few years rapidly extended its distribution from Africa and the Middle East into Europe, Russia, and across Asia. LSDV is believed to be primarily spread mechanically by blood-feeding arthropods, however the exact mode of arthropod transmission, the relative ability of different arthropod species to acquire and retain the virus, as well as their comparative importance for LSDV transmission, remain poorly characterised. Since the vector-borne nature of LSDV transmission is believed to have enabled the rapid geographic expansion of this virus, the lack of quantitative evidence on LSDV transmission has impeded effective control of the disease during the current epidemic. Obtaining high quality data on virus transmission by arthropods is challenging, and practical limitations often result in inadequate arthropod numbers or model hosts, limiting the transferability of experimental findings to the natural transmission scenario.

We have addressed these limitations in this study. Using a highly representative bovine experimental model of lumpy skin disease we allowed four representative vector species (*Aedes aegypti, Culex quinquefasciatus, Stomoxys calcitrans* and *Culicoides nubeculosus*) to blood-feed on LSDV-inoculated cattle in order to examine the acquisition and retention of LSDV by these species in unprecedented detail. We found the probability of LSDV transmission from clinical cattle to vector correlated with disease severity. Subclinical disease was more common than clinical disease in the inoculated cattle, however the probability of vectors acquiring LSDV from subclinical animals was very low.

All four potential vector species studied had a similar rate of acquisition of LSDV after feeding on the host, but *Aedes aegypti* and *Stomoxys calcitrans* retained the virus for a longer time, up to 8 days. There was no evidence of virus replication in the vector, consistent with mechanical rather than biological transmission. The parameters obtained in the in-vivo transmission experiments subsequently enabled enhanced modelling approaches to determine the basic reproduction number of LSDV in cattle mediated by each of the insect species. This was highest for *Stomoxys calcitrans* (19.1), *C. nubeculosus* (7.4), and *Ae. aegypti* (2.4), surprisingly indicating these three species are all potentially efficient transmitters of LSDV. These results reveal that currently applied LSDV control measures such as stamping out of all cattle on affected premises or insect control measures targeting single species need to be urgently reconsidered. Overall our studies have highlighted that the combination of highly relevant *in-vivo* experiments and mathematical modelling can be directly applied to devise evidence-based proportionate and targeted control programmes.

## Introduction

Lumpy skin disease virus (LSDV) is a large DNA virus of the family *Poxviridae* and the etiological agent for lumpy skin disease (LSD) in cattle. LSDV is a rapidly emerging pathogen that is believed to be mechanically transmitted by arthropod vectors. First described in Zambia in cattle in 1929, LSDV subsequently spread throughout Africa and into the Middle East [1]. In the past decade the virus has increased its geographical coverage substantially, entering and spreading within Europe and Asia including Russia, India, Bangladesh, Taiwan and China [2-8]. LSD is characterised by fever, weight loss, and prominent multifocal necrotising cutaneous lesions [9], and affects cattle of all ages [10]. Morbidity in disease outbreaks ranges from 5-26%, and mortality 0.03-2% [2-4, 11-13). Control measures include vaccination, quarantine and partial or complete culling of infected herds. The direct impact of LSD outbreaks and subsequent control measures cause significant negative economic and welfare impacts, resulting in food insecurity for affected communities in endemic [14-16] and epidemic [17] situations.

To date the mode of LSDV arthropod transmission has been assumed to be mechanical as no evidence of active virus replication in insects or ticks has been found [30]. This mechanical arthropod-borne spreads is believed to have enabled the rapid geographic expansion of LSDV, however fundamental yet crucial questions such as the species of arthropods responsible and the infectious period of LSDV-infected cattle remain unknown. This incomplete knowledge of LSDV transmission has impeded the implementation of targeted and evidence-based control measures.

Haematophagus dipterans (referred to in this work as “blood-feeding insects”), particularly *Stomoxys calcitrans*, have been associated with outbreaks of LSDV [7, 18-20], mainly based on inference of transmission patterns and insect ecological parameters. In addition, proof-of-principle experimental transmission of LSDV from affected to naïve animals (defined by the presence of clinical disease and/or detection of systemic LSDV antigen and/or capripoxvirus-specific antibodies) has being demonstrated via the mosquito *Aedes aegypti* [21], the ticks *Rhipicephalus appendiculatus* [22-24], *Rhipicephalus decoloratus* [25], *Amblyomma hebraeum* [26], the stable fly *Stomoxys calcitrans*, horseflies *Haematopota* spp. and other *Stomoxys* species [27, 28]. LSDV DNA has also been detected in other species after feeding on infected cattle or an infectious blood meal (*Culex quinquefasciatus, Anopheles stephensi, Culicoides nubeculosus*) [29], or in field-caught pools (*Culicoides punctatus*) [4]. However transmission of LSDV to susceptible animals has not been confirmed for these species.

Overall these studies provide growing evidence of the potential participation of different arthropods in the transmission of the LSDV. Unfortunately previous studies had design limitations including reduced number of donor cattle and limited times post-infection, the use of virus-spiked blood meals and/or reduced number of insects assayed. As a consequence the results obtained do not fulfil quantitative requirements to assess the risk of transmission. This is demonstrated by large uncertainty in parameter estimates. Furthermore the vital knowledge gap of understanding how efficient each vector is contributing epidemiologically to the transmission of LSDV remains.

LSDV can be detected in skin lesions, blood (primarily in peripheral blood mononuclear cells), and in nasal, oral and ocular secretions of infected cattle [27, 31, 32]. Viraemia is considered of short duration and relatively low level, though the virus can survive for longer periods of time in skin lesions [31]. LSDV has also been detected in seminal fluid of diseased bulls [33], making venereal transmission a possibility [34-36]. Subclinical infections (detection of LSDV in animals without cutaneous lesions) [3, 27, 32] and resistance to LSDV (absence of LSDV and cutaneous lesions following experimental challenge) have been reported, but both are poorly documented. The contribution of subclinical LSD to the transmission of the virus is unclear and a topic of controversy when implementing control measures such as whole-herd culling (including asymptomatic animals), particularly when morbidity is low [37, 38].

In this study we used a highly relevant experimental LSD infection model in the natural cattle host and four representative blood-feeding insect species previously reported to be capable of acquiring LSDV (*S. calcitrans, Ae. aegypti, Cx. quinquefasciatus* and *C. nubeculosus*). We obtained quantitative data describing critical, biologically relevant parameters of the mechanical transmission of LSDV in unprecedented detail. These transmission parameters subsequently enabled advanced mathematical modelling to understand the risk of transmission of the virus from experimentally infected cattle to each vector insect species. Furthermore our in-vivo transmission studies provided much-needed evidence that subclinically infected cattle do not contribute efficiently to virus acquisition by blood-feeding insect, as well as further defining the infectious period of the cattle host. In combination with published data on insect ecology parameters and feasibility of LSDV transmission our results ultimately allowed the determination of the basic reproduction number for each insect vector species. These studies highlight the powerful combination of natural host infection/ transmission studies and subsequent mathematical modelling to enable maximum relevance and transferability of data to the in-field situation and direct applicability to improve control measures.

## Results

### Experimental infection of calves with LSDV

#### Experimental inoculation of calves with LSDV results in clinical and subclinical disease

Eight calves were challenged by intravenous and intradermal inoculation of LSDV in order to act as donors on which blood-feeding insects could feed. The clinical and pathological findings have been described previously [9], and resemble those of naturally infected cattle [2, 4, 8, 11, 12, 37]. Three calves (calves 3, 5, and 9) developed lumpy skin disease, characterised by severe multifocal dermatitis with necrotising fibrinoid vasculitis consistent with field reports of LSD (S1A Fig). The cutaneous lesions initially appeared in close proximity to the inoculation site at 5 days post challenge (dpc) for calves 5 and 9, and at distant sites in all three clinical calves at 7 dpc. The five remaining calves (calves 2, 4, 7, 8 and 10) did not develop lesions other than at the inoculation sites (S1B Fig). All eight inoculated calves developed a fever which was more prolonged in calves with clinical signs (S1C Fig). Superficial lymph nodes, predominantly the superficial cervical lymph node, were enlarged in both groups starting between 2-5 dpc, with larger lymph nodes present in clinical compared to subclinical calves (S1D Fig). Two non-inoculated in-contact calves (calves 1 and 6) were included in the study and did not develop any clinical signs or lesions consistent with LSD.

#### LSDV DNA can be detected in blood and skin of clinical and subclinical calves

In the three clinically-affected calves viral DNA was first detected in the blood by qPCR at 5 dpc and remained detectable in all subsequent blood samples (up to 19 dpc). Peak viral DNA levels in the blood (6.9, 5.3 and 5.3 log_10_ copies/ml in calves 3, 5 and 9, respectively) were reached at 11 dpc (Fig 1). By contrast, viral DNA was detected only intermittently in the blood of four (out of five) subclinically infected calves between 5 dpc and 19 dpc. In addition, genome copy numbers were lower (median: 2.1 log_10_ copies/ml; range: 1.2 to 2.4 log_10_ copies/ml) than those in clinically-affected calves (Fig 1). Although negative for LSDV in whole blood, the peripheral blood mononuclear cell (PBMC) fraction of calf 7 was positive for viral DNA on days 7, 9 and 19 post challenge (Data S1). These results indicate that clinical calves had had more viral DNA present in the blood, and for longer compared to subclinical calves. However, LSDV DNA could be detected at least once in all eight challenged animals between 5 and 19 dpc.

**Fig 1.**
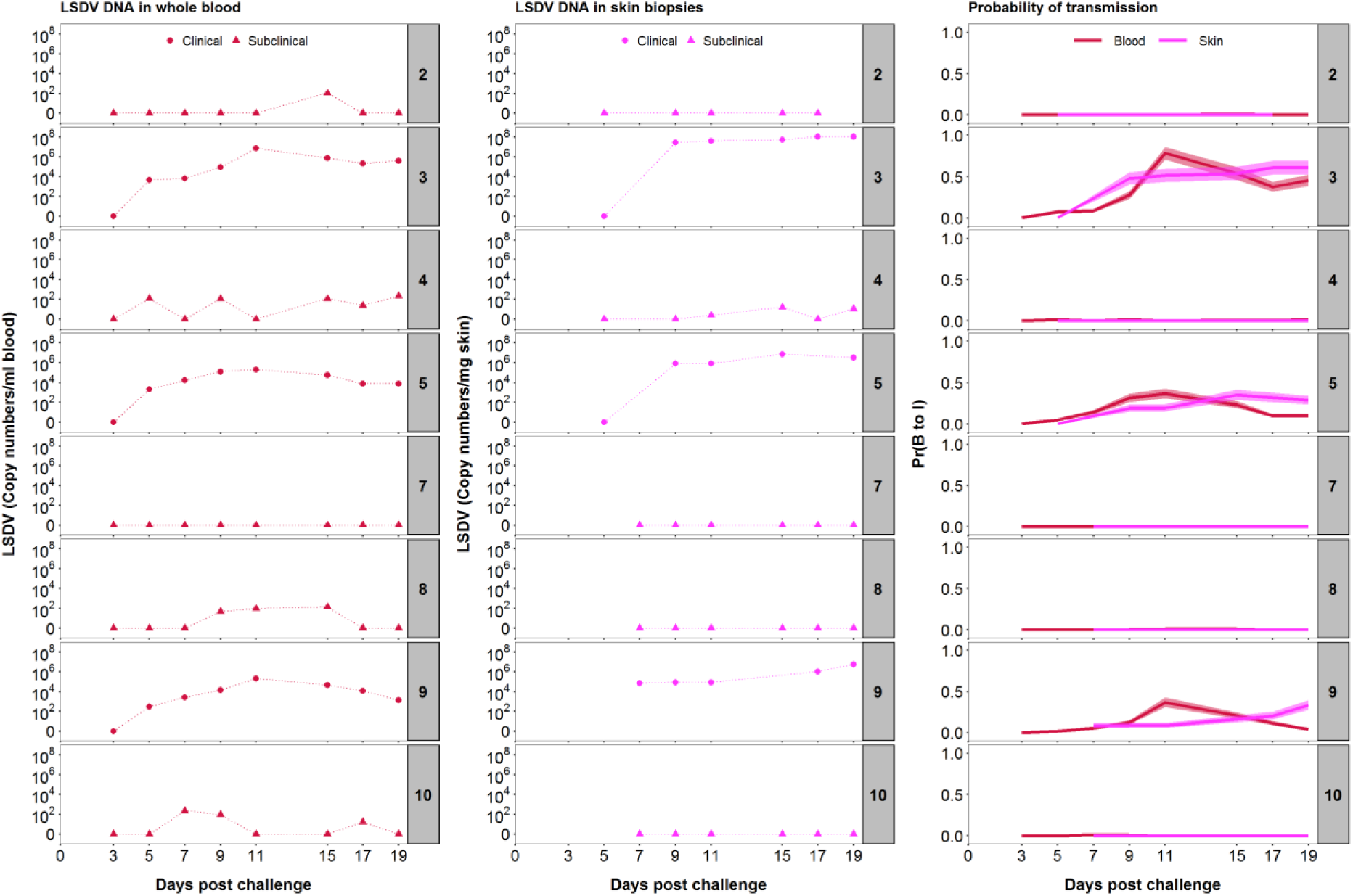
LSDV inoculation of eight calves results in a spectrum of infectiousness. Levels of viral DNA in blood (log_10_ copies/ml; first column) and skin (log_10_ copies/mg; second column) of the inoculated calves at different days post-challenge were quantified by qPCR. Based on the viral DNA levels in blood (red) or skin (magenta) the corresponding probability of transmission from bovine to insect (“infectiousness”) was calculated using a dose-response relationship (third column). Lines and shading show the posterior median and 95% credible intervals for the probability, respectively.

Skin biopsies of cutaneous lesions taken at 7 dpc (calf 9) or 9 dpc (calves 3 and 5) contained abundant viral genomes as measured by qPCR (Fig 1). Viral DNA was detected in all subsequent biopsy samples, with the quantities detected remaining at an approximately constant level for the duration of the experiment (Fig 1). The amount of viral DNA present in the skin lesions varied between the three clinical calves in an analogous fashion to the viral DNA in blood, with the highest concentration of viral DNA detected in skin lesions of calf 3 and least in calf 9 (Fig 1). The peak level of viral DNA in skin was reached after the peak level of viral DNA in blood in all three calves (Fig 1). Viral DNA was detected at three time points in biopsies of normal skin from one subclinical calf (calf 4) at a lower copy number than in the clinically-affected animals; skin biopsies from the other subclinical animals (calves 2, 7, 8 and 10) were all negative for LSDV DNA (Fig 1).

#### Infectious LSDV is present in larger quantities in the skin compared to blood

Both skin homogenate and PBMC suspension collected between 5 and 19 dpc from clinical calves were titrated to determine the quantity of live virus in these tissues. Although units of measurement are not directly comparable between sample types (i.e. skin vs PBMC), they are representative of the magnitude of exposure that haematophagus insects may encounter during feeding (i.e. mg of skin tissue and µl of blood). In all calves the viral titre from skin homogenate was higher and more constant than from PBMC suspension (Fig 2). Live virus was detected for six consecutive days from 5 dpc in the PBMC fraction of calf 3, whereas in calves 5 and 9 the virus was isolated only in three and two days (respectively) starting at day 7 post challenge. In contrast, all skin samples except one taken from dermal lesions contained live LSDV with a maximum titre of 10^4.3^ PFU/mg skin, which is over 10^3^-fold greater than the maximum level of virus detected in PBMCs, emphasising the strong cutaneous tropism of LSDV. Biopsies collected from normal skin of clinical calves were negative for live virus (i.e. below 10^−2^ PFU/mg, Data S1) suggesting the virus is highly concentrated in the skin lesions of clinical animals. Live virus was not detected in blood or skin from subclinical animals (including samples which were qPCR positive).

**Fig 2.**
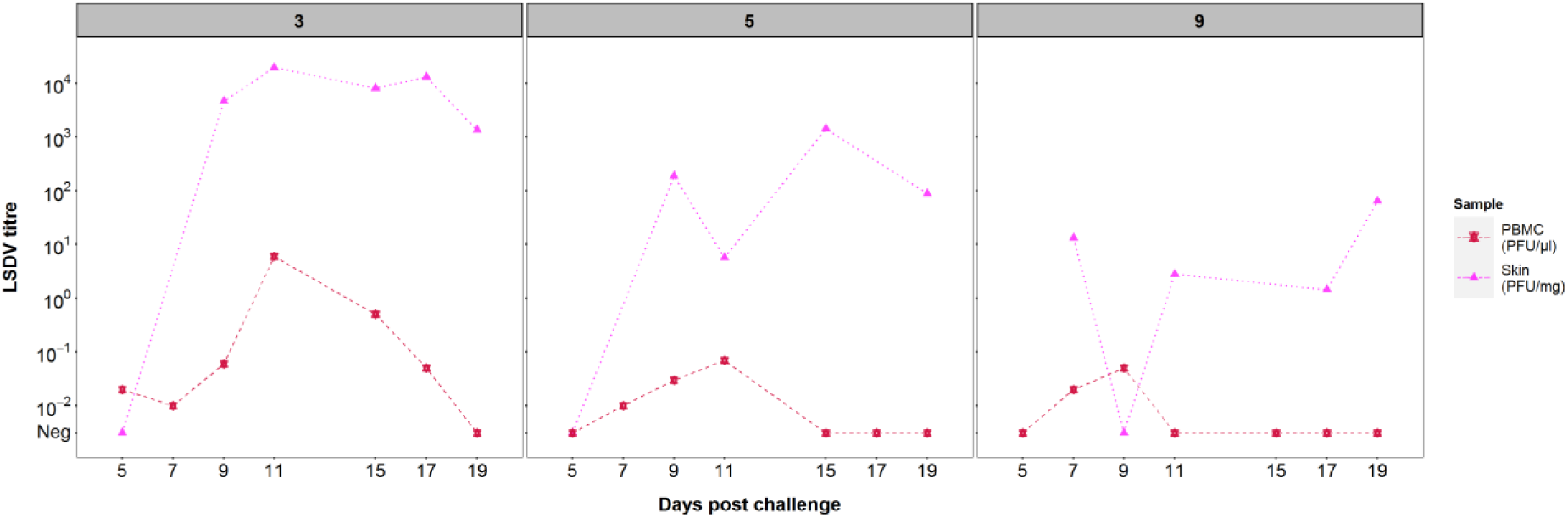
LSDV titres vary between three clinical animals but are consistently higher in the skin compared to blood. Levels of infectious lumpy skin disease virus (LSDV) in skin biopsies (PFU/mg of skin) (magenta triangles) and peripheral blood mononuclear cell (PBMC) fractions (PFU/µl suspension) (red stars) were quantified by titration on MDBK cells. Generalised skin lesions were first noted in all three animals at 7 days post challenge.

#### Humoral response to LSDV inoculation

Serum from the three clinically-affected calves contained antibodies to LSDV at 15-17 dpc as determined by a commercial ELISA test. By the end of the study period all subclinical animals had also developed detectable LSDV antibodies at levels lower than those observed in the clinical animals, but above those of the non-challenged controls (S1E Fig). The presence of detectable levels of antibodies confirmed exposure to the virus in all eight challenged animals, although the clinical outcome of challenge varied widely between the eight calves.

### Acquisition and retention of LSDV by blood-feeding insects after feeding on donor cattle

We next studied the influence of this disease spectrum on the acquisition and retention of LSDV in blood-feeding insects. To assess the acquisition and retention of LSDV by blood-feeding insects, all eight challenged animals were exposed to two mosquito species, *Ae. aegypti* and *Cx. quinquefasciatus*, one species of biting midge, *C. nubeculosus*, and the stable fly, *S. calcitrans* on days 5, 7, 9, 11, 15, 17 and 19 post challenge. The selected species are potential mechanical vectors with different feeding mechanisms [39], covering those which will feed readily on cattle (*i*.*e. S. calcitrans*), as well as species models for Culex and Aedes mosquitoes [40, 41] and also biting midges [42, 43] which would feed on cattle. At each time point, a pot of insects of each species (i.e. four pots in total) was placed on a separate cutaneous nodule on a clinical animal and, a corresponding area of normal skin on a subclinical animal. Blood engorgement, as a measure for detection of insect biting activity, was assessed visually. A subset of the insects from each pot was tested for the presence of LSDV DNA by qPCR at 0, 1, 2, 4 and 8 days post feeding (dpf) (S2 Fig). The smaller numbers of insects tested at the later time points reflect the lower numbers surviving for long enough to be tested.

Different models for the proportion of positive insects were compared to assess differences in: (i) the probability of transmission from bovine to insect (i.e. of acquiring LSDV) amongst insect species and between clinical and subclinical donors; and (ii) the duration of viral retention amongst insect species (S1 Table). Models were compared using the deviance information criterion (DIC), with a model having a lower DIC preferred to one with a higher DIC. Positive insects were those with LSDV DNA amplification by qPCR.

#### Probability of transmission from bovine to insect

A total of 3178 insects were fed on the eight donor calves (over 7 feeding sessions), of which 180 were positive for viral DNA when tested. A higher proportion of insects were positive after feeding on a clinical donor (173 out of 1159) compared to feeding on a subclinical donor (7 out of 2019) (S2 Fig). Comparing the proportion of positive insects for each species after feeding in clinical and subclinical calves (Fig 3) revealed that the probability of transmission from bovine to insect (i.e. of acquiring LSDV) does not differ amongst the four insect species, but that this probability does differ between clinical and subclinical donors (S1 Table). For a clinical donor, the probability of transmission from bovine to insect was estimated (posterior median) to be 0.22, while for a subclinical donor it was estimated to be 0.006 (Table 1). This means that an insect feeding on a subclinical animal is 97% less likely to acquire LSDV than an insect feeding on a clinical one (Table 1; Fig 3).

**Fig 3.**
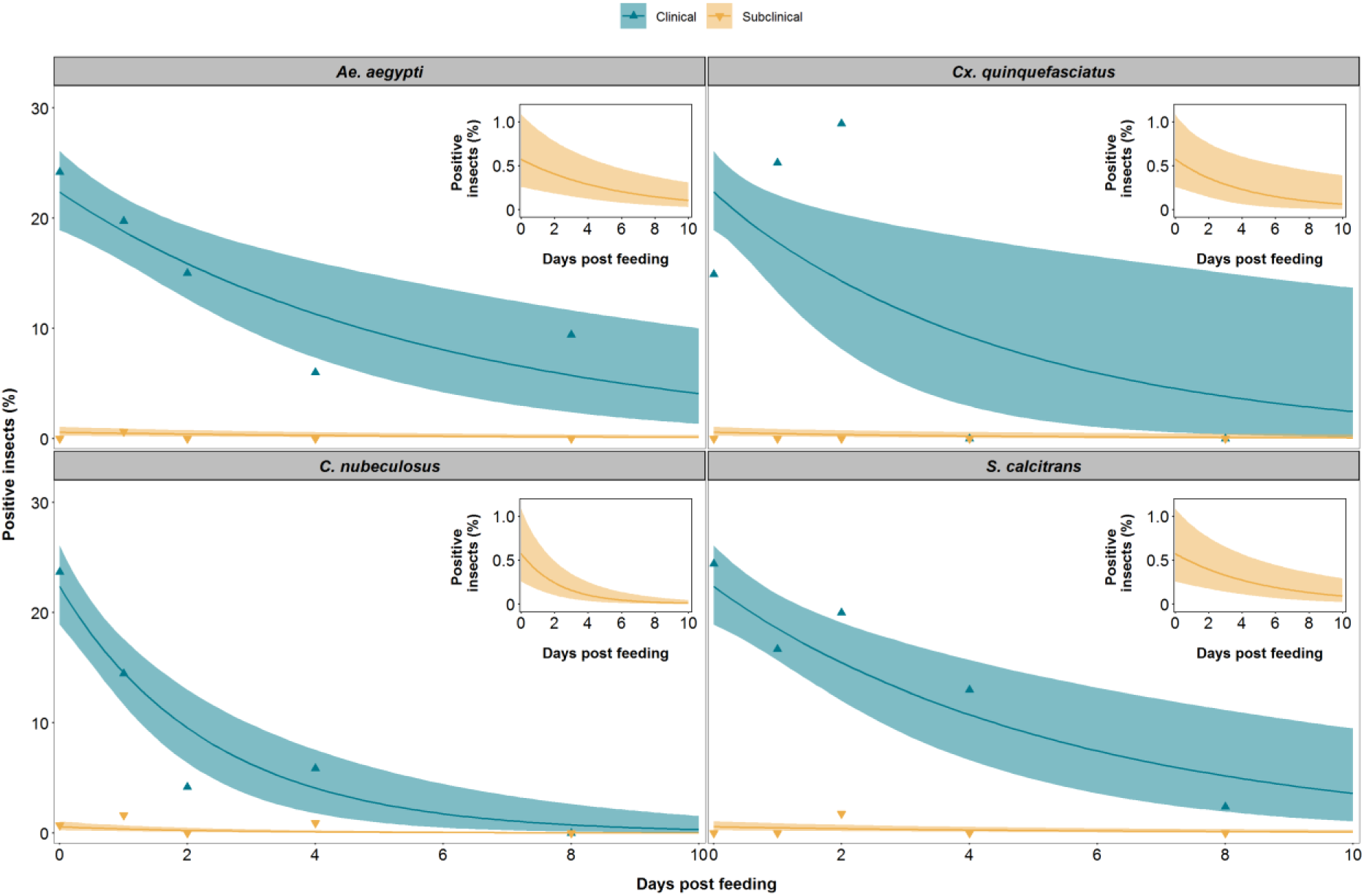
LSDV is retained in blood-feeding insects for up to 8 days post feeding. The proportion of blood-feeding insects positive for lumpy skin disease viral DNA after feeding on a clinical (green) or subclinically (yellow) animal is shown for the four species of insect: *Aedes aegypti*; *Culex quinquefasciatus*; *Culicoides nubeculosus*; and *Stomoxys calcitrans*. Each plot shows the observed proportion of positive insects (triangles) and the expected proportion of positive insects (posterior median (line), and 2.5th and 97.5th percentiles of the posterior distribution (shading)). The inset shows the expected proportion of positive insects after feeding on a subclinical animal using a graph with an expanded y-axis..

#### Infectiousness correlates with the level of viral DNA in blood and skin

The relationship between the level of viral DNA in the skin or blood of a calf and the proportion of virus-positive insects resulting from a feeding session was examined. For each feeding session that took place on the three clinical calves, the proportion of insects containing viral DNA post-feeding was calculated and compared to the viral DNA copy number present in both the blood sample and the skin biopsy taken from the calf on that day (Fig 4). This revealed a dose-response relationship between the levels of viral DNA in skin and blood and the probability of transmission from bovine to insect (or “donor infectiousness”). Furthermore, this relationship was the same for all four insect species (S1 Table), irrespective of their different feeding mechanisms. The relationship differed between levels of viral DNA in blood and skin (Table 2; Fig 4), with the probability of transmission being higher when the level of viral DNA in blood was used compared to skin (Fig 4). The fits of the models using levels of viral DNA in blood or skin are similar, suggesting that both are acceptable proxy measures for infectiousness of the donor.

**Fig 4.**
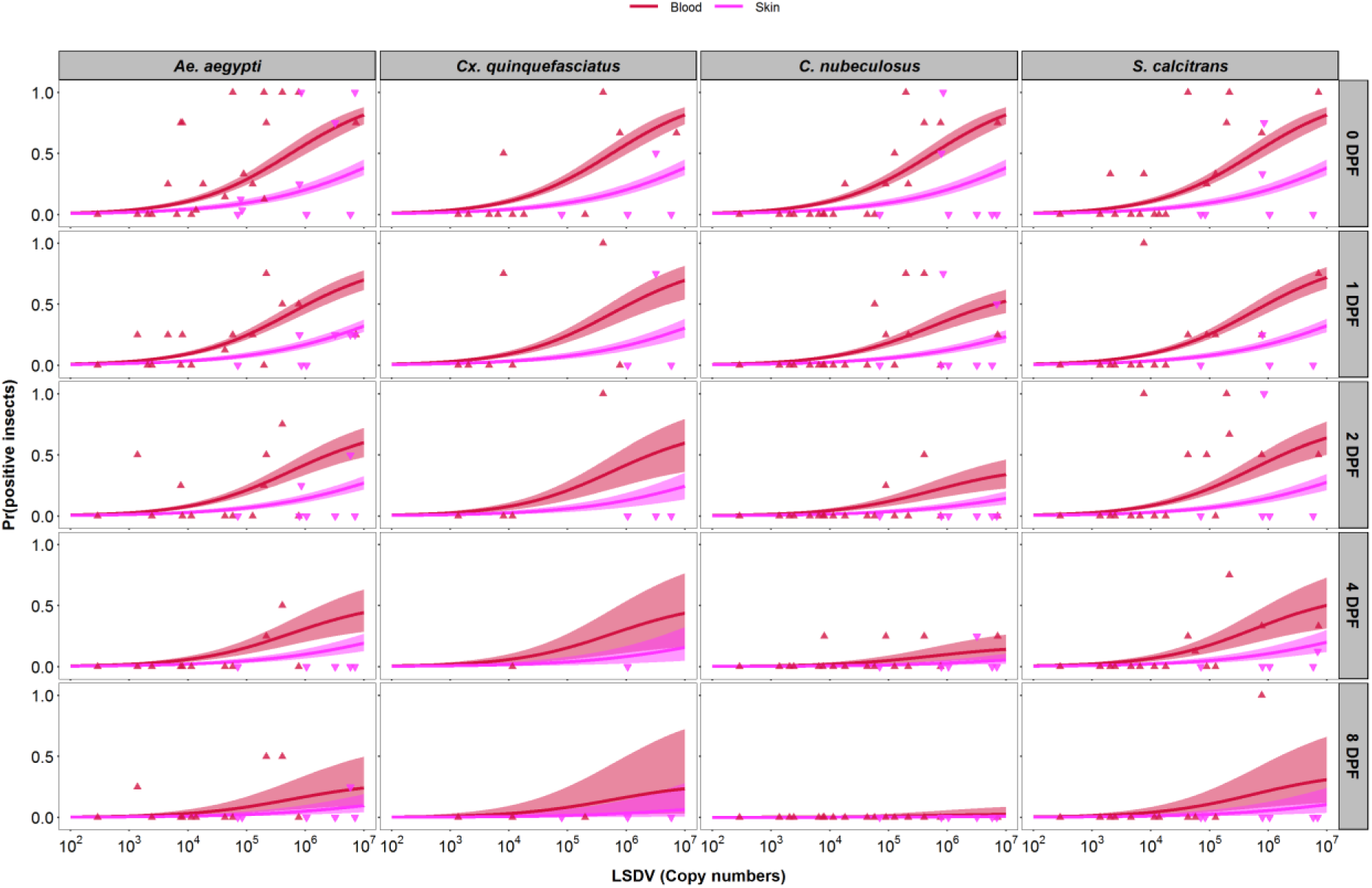
Levels of lumpy skin disease viral DNA in blood or skin are proxy measures of infectiousness. Each plot shows the dose-response relationship between the probability of an insect being positive for lumpy skin disease virus (LSDV) DNA and the level of viral DNA in the blood (log_10_ copies/ml; red) or skin (log_10_ copies/mg; magenta) of the calf on which they fed. Four species of insect, *Aedes aegypti* (first column), *Culex quinquefasciatus* (second column), *Culicoides nubeculosus* (third column) or *Stomoxys calcitrans* (fourth column) were tested at 0, 1, 2, 4 and 8 days post feeding (rows). Plots show the observed proportion of positive insects (blood: red up triangles; skin: magenta down triangles) and the estimated probability of an insect being positive (posterior median (line) and 2.5th and 97.5th percentiles of the posterior distribution (shading: blood, red; skin: magenta)).

Combining the dose-response relationship (Fig 4) with the time course for levels of viral DNA in blood or skin for each calf (Fig 1) shows how the infectiousness of an animal changes over time and how it varies amongst animals (Fig 1, right-hand column). This highlights the very low probability of transmission from bovine to insect (<0.01 at all time points; cf. estimate in Table 1) for calves which were only subclinically infected. In addition, for those calves which did develop clinical signs, there were differences in both the timing and level of infectiousness amongst the calves, which is a consequence of the underlying differences in viral dynamics in each animal. This is reflected in both the changes over time in the proportion of insects acquiring virus after feeding and differences in this proportion amongst clinical calves (S3 Fig).

#### Duration of LSDV retention

Viral DNA was detected in *Ae. aegypti* and *S. calcitrans* up to 8 dpf, in *C. nubeculosus* up to 4 dpf and in *Cx. quinquefasciatus* up to 2 dpf (Fig 3). However, few *Cx. quinquefasciatus* mosquitoes survived to 4 or 8 dpf (S2 Fig), resulting in uncertainty about the duration of retention in this species (Figs 3 and 4). The mean duration of viral retention differed amongst the four insect species in the present study (Fig 3; S1 Table), being the longest for *Ae. aegypti* (5.9 days) and *S. calcitrans* (5.5 days), followed by *Cx. quinquefasciatus* (4.5 days), and *C. nubeculosus* (2.4 days) (Fig 3; Table 1). The corresponding virus inactivation rate (i.e. the reciprocal of the mean duration of retention) was 0.17/day for *Ae. aegypti* and 0.18/day for *S. calcitrans*, 0.22/day for *Cx. quinquefasciatus* and 0.42/day for *C. nubeculosus* (Table 1).

#### Levels of retained LSDV

The median amount of viral DNA in homogenized whole insects was the same when tested at different days post feeding for three (out of the four) species: *Ae. aegypti* (Kruskal-Wallis test: *χ*^2^=0.98, df=4, *P*=0.91), *Cx. quinquefasciatus* (Kruskal-Wallis test: *χ*^2^=3.62, df=2, *P*=0.16) or *S. calcitrans* (Kruskal-Wallis test: *χ*^2^=2.74, df=4, *P*=0.60) (S3 Fig). However, the median level of viral DNA was lower for individual *C. nubeculosus* tested at later times post feeding (Kruskal-Wallis test: *χ*^2^=10.8, df=3, *P*=0.01) (S4 Fig). These results are consistent with a mechanical rather than a biological form of vector-transmission.

### Probability of transmission from insect to bovine

Three previous studies have investigated the transmission of LSDV from insects to cattle, where insects of species included in the present study were allowed to feed on an infected donor and were subsequently allowed to refeed on a naïve recipient [21, 27, 29]. The number of positive insects refeeding was not determined in these studies. By combining LSDV acquisition and retention results of the present study with challenge outcomes of the aforementioned studies (i.e. whether or not transmission occurred), it is possible to estimate the probability of transmission from insect to bovine. This probability was highest for *Ae. aegypti* (0.56), intermediate for *C. nubeculosus* (0.19) and *Cx. quinquefasciatus* (0.11) and lowest for *S. calcitrans* (0.05) (Table 1). However, there is considerable uncertainty in the estimates for all species, but especially for *Ae. aegypti, C. nubeculosus* and *Cx. quinquefasciatus* (Table 1), which makes it difficult to compare estimates across species.

### Basic reproduction number for LSDV

The basic reproduction number (*R*_0_) is defined as “the average number of secondary cases caused by an average primary case in an entirely susceptible population” [44]. For LSDV, *R*_0_ combines the parameters related to transmission (Table 1) with those related to vector life history (i.e. biting rate, vector to host ratio and vector mortality rate; see Table 1 in ref. 45) to provide an overall picture of the risk of transmission by the four insect species [45]. The basic reproduction number was estimated to be highest for *S. calcitrans* (median *R*_0_=19.1) (Table 1; Fig 5), indicating that this species is likely to be the most efficient vector of LSDV and would be able to cause substantial outbreaks if it were the sole vector in a region. Both *C. nubeculosus* (median *R*_0_=7.1) and *Ae. aegypti* (median *R*_0_=2.4) are also potentially efficient vectors of LSDV (i.e. *R*_0_>1 for these species) and would be able to sustain transmission if either were the sole vector in a region. Finally, *Cx. quinquefasciatus* (median *R*_0_=0.6) is likely to be inefficient at transmitting LSDV (Table 1; Fig 5). It would not be able to sustain transmission on its own, but it could contribute to transmission if other vector species were also present.

**Fig 5.**
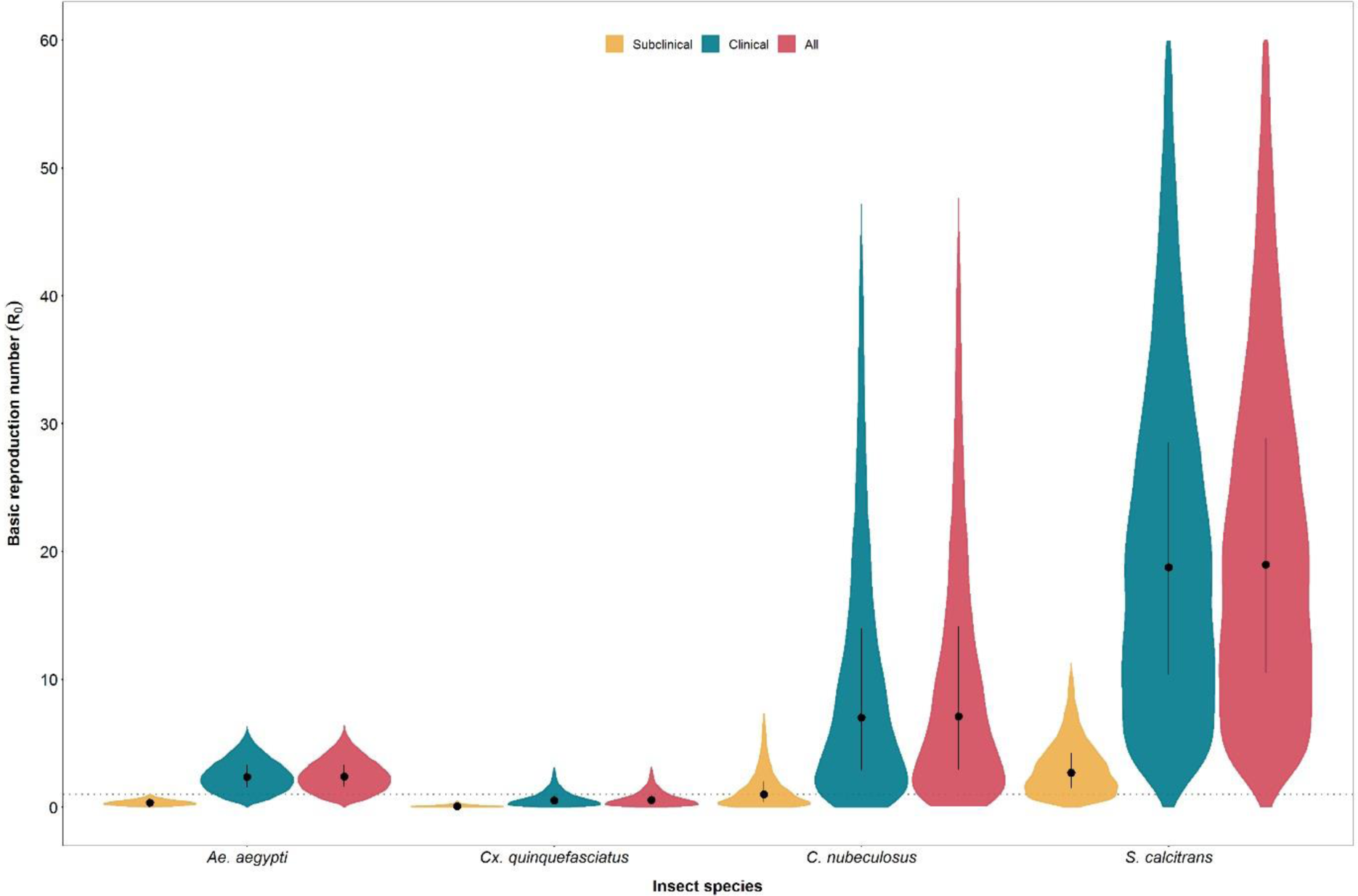
Basic reproduction number (*R*_0_) for lumpy skin disease virus (LSDV) in calves when transmitted by *Aedes aegypti, Culex quinquefasciatus, Culicoides nubeculosus* or *Stomoxys calcitrans*. For each species, *R*_0_ was calculated for subclinical calves only (yellow), clinical calves only (green) and both combined (red). Violin plots show the posterior median (black circle), interquartile range (black vertical line) and density (shape) for *R*_0_ based on replicated Latin hypercube sampling (100 replicates with the range for each parameter subdivided into 100 steps).

Exploring the contribution of clinical and subclinical animals to the basic reproduction number for each species further emphasises the more limited role played by subclinical animals in the transmission of LSDV (Fig 5). For all species, the *R*_0_ for clinical animals alone is very close to that for both clinical and subclinical animals combined (Fig 5). Moreover, the median *R*_0_ for subclinical animals alone is below one for all species, except *S. calcitrans* (Fig 5).

The *R*_0_ values calculated from our data and previous studies provide a summary of the risk of LSDV transmission. A range of blood-feeding insects are likely to support a disease outbreak by transmitting LSDV from a clinical to a naïve animal, particularly biting flies such as *S. calcitrans*. The *R*_0_ calculations also highlight that, although there may be a significant subset of subclinical animals in an affected herd, they are likely to play at most a minor role in the transmission of the virus.

## Discussion

This study describes a controlled experimental model of LSD that mimics disease features described in field outbreaks [2, 4, 8, 11, 12, 37] and other experimental models [27, 32]. Inoculated calves (both clinical and subclinical) were used to measure the acquisition (transmission from bovine to insect) and retention of LSDV by four potential vector species. These data were then used to estimate the risk of transmission by these species with the aim of providing evidence with which to inform decisions during the implementation of measures to control LSDV.

In our experimental model we observed that 37.5% of calves developed generalised LSD with the remaining 62.5% of calves classified as subclinical (no cutaneous nodules, positive qPCR in blood [27]). This attack rate of 0.37 is comparable to other experimental models with field strains of LSDV (0.57 [27] and 0.50 [32]). Reports of animals with subclinical LSD in the field is sparse, with an incidence of up to 31.3% reported [3]. The high detection of subclinical infection in our study may be a result of an intense sampling protocol (compared to the limited sampling of individuals during an outbreak investigation). Further investigation of the true incidence of subclinical LSD in field studies is warranted.

Cattle experimentally infected with LSDV, including in our study, have higher concentrations of LSDV in skin lesions than blood (Figs 1 and 2). In clinically infected animals we identified a relationship between the viral load in skin and blood and the proportion of insects positive for the virus, indicating both skin and blood are good predictors of the transmissibility of LSDV from donors to vector. Our study did not extend beyond 21 days post challenge however, and this observation may only be true during the initial stage of the disease when the viraemia is detectable. Donors with different disease severity and therefore different levels of infectiousness would strongly influence the proportion of vectors which acquired virus. This finding may explain the discrepancies between experimental studies which have assessed the transmission of LSDV by vectors [21, 29] when the infectiousness of the donors may have been different.

As reported in this study and others [27, 31] LSDV can be detected in the blood of cattle prior to the appearance of skin lesions, 5-8 dpc. However, during this time, viraemia is relatively low and in our study few insects were positive for LSDV after feeding (S3 Fig). Viraemia rises and peaks after the multifocal skin lesions appear (at around 7 dpc), and this is when the probability of transmission from bovine to insect starts to increase (Fig 1). The probability remains high while viraemia is high and when skin lesions are present. The appearance of skin lesions therefore marks the start of the risk period for virus transmission, and this means that rapid diagnosis and consequent implementation of control measures should be possible and effective at limiting onwards transmission [46, 47]. In this study we were only able to follow the animals for 21 days post-challenge with the last exposure of blood-feeding insects to infected calves on day 19, and thus the period for transmission risk could not be established beyond this time point. Nevertheless, under controlled conditions [31], LSDV has been isolated up to 28 (blood) and 39 (skin) days post-challenge, and detected by PCR up to 91 days post-challenge (in skin biopsies). Therefore, LSDV uptake by vectors may occur beyond the reported period in our study.

We found that subclinical donors were much less likely than clinical animals to transmit virus to vectors (Table 1; Fig 3), indicating a substantially reduced role of subclinically infected animals in the transmission of LSDV. For some vector-borne diseases such as dengue fever, malaria, asymptomatic and preclinical individuals may be an important source of the pathogen for vectors and help maintain the transmission cycle [48, 49]. The situation with LSDV appears to be different. The viraemia in subclinical animals is low and skin lesions (representing the major viral load) are absent in these animals. Few vectors therefore acquire LSDV from subclinical cattle, and this reduces the chances of onward transmission to a susceptible host. This is the first time the relative contribution of subclinically infected cattle to onward transmission of LSDV has been quantified.

Lumpy skin disease virus can be mechanically transmitted by stable and horse flies [27, 28] and mosquitoes [21]. Mechanical transmission of viruses by blood-feeding vectors can be influenced by their feeding mechanism, ecology and biting behaviour. Stable flies are aggressive feeders with a painful bite which leads to interrupted feeding and to more than one feeding event per day [50, 51]. They are also known to regurgitate previous blood intakes while feeding. To penetrate the skin, stable flies rotate sharp teeth on their proboscis (5-8mm long) and form a pool of blood from which they feed [39]. *Culicoides* midges also disrupt the skin barrier using their proboscis (0.1-0.2 mm long). Midges serrate the skin using saw-like blades on their proboscis that cross over each other to produce a pool of blood [39]. Biting midges feed generally less frequently than stable flies as feeding is associating to their gonotrophic cycle (7-10 days, but as a temperature-dependent event it can be as short as 2-3 days) [52]. Mosquitoes do not produce pools of blood, instead they penetrate the skin “surgically” searching for a capillary with their proboscis of (1.5-2.0 mm long), accompanied by a pushing and withdrawing movement until it hits a capillary from which to withdraw blood [53]. Mosquitoes feeding on blood is also associated to their gonotrophic cycle, but multiple feedings have been reported in some species [54, 55] Despite these variations in feeding behaviour, all four insect species acquire LSDV at the same rate, indicating that virus acquisition is not influenced by feeding behaviour.

All four insect species in the present study were able to acquire LSDV through feeding on clinical animals and to retain it for several days (Fig 3). In a small proportion of *Ae. aegypti* and *S. calcitrans* LSDV DNA was still present at 8 days post feeding, which was the longest we investigated, thus longer retention cannot be ruled out. Similar to our study, Chihota and co-authors [21], identified that *Ae. aegypti* mosquitoes feeding on animals with clinical LSD were able to acquire and retained the virus for up to six days, and that the proportion of virus-positive insects also decreased with days post feeding. They observed similar dynamics in *Cx. quinquefasciatus* and *Anopheles stephensi* mosquitoes when using a membrane-feeding system with a LSDV-infected bloodmeal, but when they fed *C. nubeculosus* and *S. calcitrans* on LSDV-infected calves they did not detect the virus beyond the day of feeding (*C. nubeculosus*) or the following day (*S. calcitrans*) [29]. However, we now know that disease severity of the donor can influence the acquisition of LSDV by an insect. This could result in the lower acquisition and retention observed by Chihota et al. In our work we identified that LSDV can be retained longer than previously reported in *S. calcitrans* and *C. nubeculosus*, with a decline in virus DNA post-feeding only detectable for *C. nubeculosus*.

For LSDV, as for other chordopoxviruses including capripoxvirus, fowlpox virus and myxoma virus, the mode of vector-mediated transmission is assumed to be mechanical [21, 56-59]. Our data and that of Chihota and co-authors support the theory that LSDV does not replicate in the insect (at least at detectable levels), but the retention of viral DNA in *Ae. aegypti* and *S. calcitrans* at levels similar to those acquired during feeding deserves further investigation [60].

Assessment of acquisition and retention of LSDV genome was performed in whole insect homogenates in our study, and further investigations into the location of virus within the insects were not possible. However an earlier study with *Ae. aegypti* [61] indicated that LSDV DNA persist longer in the head than in the thorax/abdomen. This is consistent with research that found myxoma virus was retained on the mouthparts of *Ae. aegypti* mosquitoes up to 28 days post feeding [62]. The mechanism by which poxviruses persist for days on the mouthparts of vectors warrants further study.

The detection of LSDV in insect vectors in our study was based on the presence of viral DNA rather than infectious virus particles. Virus titration from homogenates of individual insects was attempted however we were able to detect live virus only from pooled homogenates of *S. calcitrans* and of *Ae. aegypti* (data not shown), suggesting low numbers of infectious virions are present on each insect. In previous work live LSDV was detected in individual *Ae. aegypti* for up to 6 days following exposure to an infectious calf [21], and live goatpox virus up to 4 days in *S. calcitrans* [59].

The aim of the present study was to use the results of feeding four model vector species on LSDV-infected cattle to estimate parameters related to transmission that were not possible with the data from previous studies. Given the large number of insects fed and tested (>3000), the resulting estimates for the probability of transmission from bovine to insect (including the relative risk of transmission from a subclinical animal and the dose-response) are robust, as indicated by the narrow credible intervals for these parameters (Tables 1 and 2). The estimates for the duration of virus retention (or, equivalently, the virus inactivation rate) are more uncertain (Table 1), which reflects difficulties in keeping insects alive to later days post feeding, especially *Cx. quinquefasciatus* (S2 Fig).

Although not assessed in the present study, we used data from previous transmission experiments [21, 27] to estimate the probability of transmission of LSDV from insect to bovine. The small number of studies (and animals in each study) mean that the estimates for this parameter are uncertain, extremely so for *Ae. aegypti, Cx. quinquefasciatus* and *C. nubeculosus* (Table 1). This uncertainty is less important for *Cx. quinquefasciatus*, which is unlikely to be an important vector even if were able to transmit LSDV efficiently, but it makes it difficult to determine whether or not *C. nubeculosus* is likely be an important vector.

One previous study assessed the importance of different vector species by calculating *R*_0_ for LSDV from its component parameters [45], based on data from two studies by Chihota and co-authors [21, 29]. Despite different values for the underlying parameters, both this and the present study obtained similar estimates of *R*_0_ for *S. calcitrans* (median: 15.5 vs 19.1) and for *Cx. quinquefasciatus* (median: 0.8 vs 0.6), suggesting the former is likely to be an important vector and the latter is likely to be an inefficient vector of LSDV. The estimates of *R*_0_ for *Ae. aegypti* differed between the studies (median: 7.4 vs 2.4), due to differences in estimates for the mean duration of virus retention (11.2 vs 5.9) and probability of transmission from bovine to insect (0.90 vs 0.22). This suggests that *Ae. aegypti* (or other *Aedes* spp. for which it could be considered a model) may be less efficient vector than previously assumed. Finally, there was a major difference between the studies in their estimates of *R*_0_ for *C. nubeculosus*. The earlier study suggested this species is likely to be a less efficient vector (median *R*_0_=1.8), but the present one suggests it could be an efficient one (median *R*_0_=7.4). This discrepancy is principally related to markedly different estimates for the mean duration of virus retention (0.01 days vs 2.4 days). Moreover, the estimate of *R*_0_ for this species is highly uncertain, largely as a consequence of uncertainty in the estimate of the probability of transmission from insect to bovine (Table 1). *Culicoides* spp. are ubiquitous on cattle farms [63, 64] and, consequently, would represent a major transmission risk if they proved to be efficient vectors of LSDV. Hence, it is important that their ability to transmit virus to cattle is assessed.

Linking transmission experiments with mathematical modelling is an uncommon and powerful approach to create robust evidence which can inform policy makers involved in controlling the spread of infectious diseases. Here we have used this approach to investigate the transmission of LSDV, which has recently emerged as a significant threat to cattle in Africa, Asia and Europe. Our evidence indicates that *S. calcitrans* is likely to be an important vector species. It also suggests that *Culicoides* biting midges may be a more efficient vector species than previously considered. Furthermore, we have demonstrated for the first time that subclinical infected cattle pose only very limited risk of onward transmission of LSDV to potential vectors. This evidence supports LSD control programmes which target clinically-affected cattle for rapid removal, rather than complete stamping-out of all cattle in an affected herd.

## Methods

### Experimental design

#### Ethical statement, housing and husbandry

The experimental study was conducted under the project license P2137C5BC from the UK Home Office according to the Animals (Scientific Procedures) Act 1986. The study was approved by the Pirbright Institute Animal Welfare and Ethical Review Board. Cattle were housed in the primary high containment animal facilities (Biosafety Level 3 Agriculture) at The Pirbright Institute. The husbandry of the animals during the study was described previously [9].

#### Challenge study and experimental procedures

Ten Holstein-Friesian male cattle (referred to as calves) were used for the study, which was done in two experimental replicates of five animals each. The median age and weight of the calves was 104 days old, 145 kg in replicate one and 124 days old, 176 kg in the second replicate. Eight calves were challenged by intravenous and intradermal inoculation with a suspension of LSDV containing 10^6^ PFU/ml [9]. More specifically, 2 ml were inoculated intravenously (jugular vein) and 1 ml was inoculated intradermally in four sites (0.25 ml in each site), two on each side of the neck.

The remaining two calves were not challenged and were kept as non-inoculated in-contact controls. Calves were randomly assigned to either the control or challenge groups using a random number generator (excluding control calf 1, which was assigned as a control on welfare grounds following diagnosis with shipping fever pneumonia). The calves were kept for 21 days following the challenge; clinical scores were taken daily and serum, whole blood and skin biopsies [9] collected over the study period. The non-steroidal anti-inflammatory drug meloxicam (0.5 mg/kg body weight) (Metacam 20 mg/ml solution, Boehringer Ingelheim) was used when required on welfare grounds.

#### Insect exposure

Blood-feeding insects used in the study were: *Aedes aegypti* ‘Liverpool’ strain, *Culex quinquefasciatus* TPRI line (Tropical Pesticides Research Institute, obtained from the London School of Hygiene and Tropical Medicine, London, UK), *Stomoxys calcitrans* (colony established in 2011 from individuals kindly provided by the Mosquito and Fly Research Unit, USDA Florida) and *Culicoides nubeculosus* [65]. All insects were reared at The Pirbright Institute under the following insectary conditions. *Ae. aegypti* were reared in pans of 300 larvae per pan, containing approximately 1 litre of water supplemented with fish food and housed at 28 °C, 70% relative humidity (RH) and 12:12 light/dark cycle. *Cx. quinquefasciatus* were reared in pans of 500-800 larvae per larval bowl, containing approximately 1.5 litre of water supplemented with ground guinea pig food and maintained at 26 °C, 50% RH and 16:8 light/dark cycle. *S. calcitrans* were reared in approximately 200 eggs per pot, incubated for 12-13 days in larval pots containing a ratio of 3:2:1 (powdered grass meal, water and corn flour) and a table spoon of yeast. *C. nubeculosus* were reared in approximately 10,000 larvae per pan containing 2 litres of dechlorinated water supplemented with oxoid broth and dried grass/wheat germ mix. Pots of 800 *Culicoides* pupae were made with males and females and allowed to emerge. Both *S. calcitrans* and *C. nubeculosus* were maintained in insectaries at 27 ± 2 °C, 50% RH, with a 16:8 light/dark cycle.

The age and sex composition of the insects at exposure was: female and male *C. nubeculosus* between 0-2 days post-eclosion, female *Cx. quinquefasciatus* and *Ae. aegypti* at 5-7 days post-eclosion and male and female *S. calcitrans* at an average of 4 days post-eclosion (range: 2-7 days). All adult insects were maintained on 10% sucrose and starved 18-24 hours before exposure to the calves.

All eight challenged calves, independent of clinical status, were exposed (for between 5 and 20 minutes) to each of the four insect species on days 5, 7, 9, 11, 15, 17 and 19 post challenge. At each time point each pot of insects was placed on a cutaneous nodule on a clinical animal and a corresponding area of normal skin on a subclinical animal. The hair of the calf at each feeding site was clipped and/or shaved, and the insects were held in close contact with the skin of the calves in a container covered by a mesh. Around two hours after exposure, insects were anaesthetised under CO_2_ and unfed individuals discarded and blood-engorged individuals collected.

For *Ae. aegypti, Cx. quinquefasciatus* and *C. nubeculosus* blood engorgement was assessed visually by the presence of blood in the abdominal cavity. However, *S. calcitrans* were all collected “blind” and blood engorgement was confirmed by the detection of the bovine cytochrome *b* gene using qPCR. Those individuals negative for cytochrome *b* at collection were removed from the analysis.

Samples from each insect group taken immediately following blood-feeding assessment (dpf 0) were stored at −80C, and the rest of the insects were maintained for 1, 2, 4 or 8 dpf. After this incubation period, surviving individuals were collected and stored at −80 °C after the incubation period. Throughout incubation all insects were maintained on 10% sucrose solution, except *S. calcitrans* which were maintained with defibrinated horse blood (TCS Biosciences Ltd) after 2 dpf. All insects were kept in a temperature-controlled room at biocontainment level 3, with a 10:14 light/dark cycle. For the incubation, cardboard/waxed pots containing the insects were placed inside plastic boxes covered by a mesh which were kept under a plastic shelter to minimise temperature and humidity fluctuations.

Temperature (mean: 24.8°C; range: 22.4°C – 26.4°C) and RH (mean: 35.9%; range: 18.5% – 48.9%) of the room and of the incubation area were recorded approximately every 15 minutes (RF513, Comark Instruments and HOBO UX100-003, Onset).

#### Samples

Skin biopsies were weighed on a calibrated scale (EP613C Explorer Pro, OHAUS®) and homogenised in 500 µl high-glucose Dulbecco’s modified Eagle’s medium (41965, Life Technologies) supplemented with 5% foetal bovine serum (Antibody Production Services Ltd, Bedford, UK), 100 U/ml penicillin and 100 μg/ml streptomycin (15140122, Life Technologies) and 2.5 μg/ml amphotericin B (15290026, Life Technologies) in a Lysing Matrix A tube (SKU 116910050-CF, MP Biomedicals) using a portable homogeniser (BeadBug Microtube Homogenizer, D1030, Benchmark Scientific Inc.). Whole insects were homogenised using a TissueLyser® (Qiagen, UK) with one or two steel beads of 3 mm (Dejay Distribution, UK) [66] in 200 µl Dulbecco’s phosphate buffered saline (PBS, 14190094, Life Technologies), supplemented with penicillin-streptomycin and amphotericin B, as above.

Bovine peripheral blood mononuclear cells (PBMC) were isolated from 7 ml of whole blood in EDTA diluted in PBS 1:1. The diluted blood was added to a SepMate™-50 centrifugation tube (Stemcell Technologies) under-layered with Histopaque®-1083 (Sigma-Aldrich). Tubes were centrifuged at 1500×g for 30 minutes, 20 °C with no brake. PBMCs were aspirated from the interface into PBS, washed three times with PBS at 1000×g for 10 minutes at 20 °C. After the final wash, cells were resuspended in 2 ml of RPMI medium (21875091, Life Technologies) supplemented with 10% foetal bovine serum, and penicillin-streptomycin as above. Blood collected without anticoagulants was allowed to clot, spun at 1000×g to 2000×g for 10 minutes in a refrigerated centrifuge and the serum collected. All samples were stored at −80 °C until analysed.

#### Laboratory assays

Nucleic acid from 200 µl of whole blood, PBMC suspension, skin homogenate or 100 µl of insect homogenate was extracted in a 96-well plate with the MagMAX™ CORE Nucleic Acid Purification Kit (Applied Biosystems, A32700) using protocol A in a KingFisher™ Flex Magnetic Particle Processor (Applied Biosystems) and eluted in 50 µl of buffer. qPCR for LSDV ORF074 detection was performed using a modification of the TaqMan assay described by Bowden *et al*. [67] with the Path-ID™ qPCR Master Mix (Life Technologies #4388644). Briefly, a 20 µl reaction was prepared using 5 µl of template, 400 nM of each primer, 250 nM of the probe and nuclease-free water to the final volume. Samples were prepared in a 96-well plate and assayed using the Applied Biosystems™ 7500 Fast Real-Time PCR System with the program: 95 °C for 10 min, and 45 cycles of 95 °C for 15 s and 60 °C for 60 s. Tissue culture derived LSDV positive controls were included in the extraction plates, and the copy number of LSDV genome were quantified using gBlocks® Gene Fragments (Integrated DNA Technologies) to generate the standard curve. The gBlocks® Gene Fragment included the target sequence of the Bowden assay for detection of LSDV ORF074 (AAATGAAACCAATGGATGGGATACATAGTAAGAAAAATCAGGAAATCTATGAGCCATCCATTTTCCAACTCTATTCCATATACCGTTTT). Bovine blood intake by insects was determined using a SYBR green assay (PowerUp™ SYBR™ Green Master Mix, A25779, Life Technologies) for the detection of bovine mitochondrial *cytochrome b* as described by Van Der Saag *et al*. [68] with some modifications (forward primer: 5’ GTAGACAAAGCAACCCTTAC at 300nM; reverse primer: 5’ GGAGGAATAGTAGGTGGAC at 500nM) using the manufacturer cycling conditions for primers with Tm >60 °C. The assay was performed in 10 µl reaction using 2 µl of template. This assay was specific for bovine *cytochrome b* and melt curve analysis was performed to confirm that only specific amplification occurred. For all qPCR assays a constant fluorescence threshold was set which produced a reproducible Cq values for the positive control samples between runs. A double antigen ELISA (ID Screen® Capripox, IDvet) was used to detect circulating antibodies for LSDV in serum samples following the manufacturer’s protocol and analysed with the Multiskan FC Microplate photometer (Thermo Scientific™). Infectious virus titrations of PBMC suspension, insect and skin homogenate was performed by viral plaque quantification in MDBK cells.

### Parameter estimation

Full details of parameter estimation are provided in S1 Text. Briefly:

#### Probability of transmission from bovine to insect and virus inactivation rate

The numbers of insects positive for viral DNA after feeding on cattle infected with LSDV were used to estimate the probability of transmission from bovine to insect and the virus inactivation rate. The probability that an insect would be positive when tested is

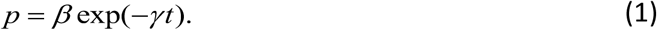

where *β* is the probability of transmission from bovine to insect, *γ* is the virus inactivation rate (i.e. the reciprocal of the mean duration of virus retention) and *t* is the time post feeding at which the insect was tested. Equation (1) combines the probability that an insect acquired virus (*β*; i.e. the probability of transmission from bovine to insect) and the probability that the insect retained the virus until it was tested at *t* days post feeding (exp(- *γt*)).

Differences amongst insect species in the virus inactivation rate and probability of transmission from bovine to insect and in the probability of transmission between subclinical and clinical animals were explored by comparing the fit of models in which these parameters did or did not vary with species or clinical status of the donor cattle. In addition, the dose-response relationship was investigated by allowing the probability of transmission from bovine to insect to depend on the level of viral DNA (in either blood or skin) in the donor animal, so that,

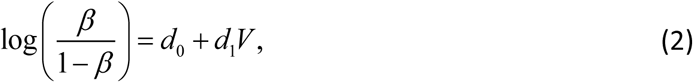

where *d*_*0*_ and *d*_*1*_ are the dose-response parameters and *V* is the level of viral DNA (log_10_ copies/ml in blood or log_10_ copies/mg in skin) in the donor when the insect fed. The different models were compared using the deviance information criterion [69]. The two proxy measures for infectiousness (i.e. level of viral DNA in blood or skin) were compared by computing posterior predictive *P*-values for each insect.

#### Probability of transmission from insect to bovine

Data on transmission of LSDV from insect to bovine were extracted from the published literature [21, 27, 29]. In these experiments, batches of insects (of the same species as used in the present study) were allowed to feed on an infected bovine and then to refeed at later time points on a naïve recipient. The probability of the recipient becoming infected is

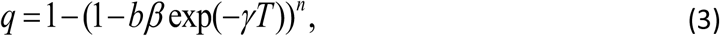

where *b* is the probability of transmission from insect to bovine, *β* is the probability of transmission from bovine to insect, *γ* is the virus inactivation rate, *T* is the time interval between feeding on the donor and refeeding on the recipient and *n* is the number of insects which refed. The probability, (3), is the probability that at least one insect (out of the *n* refeeding) transmitted LSDV, where the probability that an individual insect will transmit is the product of the probabilities that it acquired the virus during the initial feed (*β*), retained it until refeeding (exp(-*γT*)) and that it subsequently transmitted LSDV at refeeding (*b*).

#### Latent and infectious periods in cattle

Previous estimates for the latent and infectious periods of LSDV [45] were updated using the data on detection of LSDV in blood and skin collected during the present and other recently published studies [27, 32]. In addition, the proportion of cattle that develop clinical disease following challenge was estimated using data extracted from the published literature [27, 31, 32, 70, 71] and the present study.

#### Bayesian methods

Parameters were estimated using Bayesian methods. For all analyses, samples from the joint posterior distribution were generated using an adaptive Metropolis scheme [72], modified so that the scaling factor was tuned during burn-in to ensure an acceptance rate of between 20% and 40% for more efficient sampling of the target distribution [73]. The adaptive Metropolis schemes were implemented in Matlab (version 2019b; The Mathworks Inc.) and the code is available online at https://github.com/SimonGubbins/LSDVAcquisitionAndRetentionByInsects. Two chains were allowed to burn-in and then run to generate an effective sample size of around 5,000 samples (assessed using the mcmcse package [74] in R (version 3.6.1 [75]). Convergence of the chains was assessed visually and using the Gelman-Rubin statistic provided in the coda package [76] in R [75]. Different models for the variation amongst species in virus inactivation and probability of transmission from bovine to insect (S1 Table) were compared using the deviance information criterion [69].

### Basic reproduction number for LSDV

The basic reproduction number, denoted by *R*_0_, is the “average number of secondary cases arising from the introduction of a single infected individual into an otherwise susceptible population” [44]. The basic reproduction number for LSDV is,

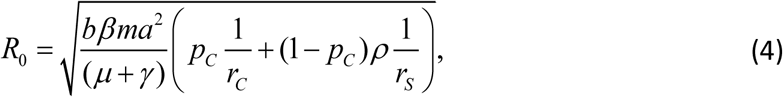

where *b* is the probability of transmission from insect to bovine, *β* is the probability of transmission from bovine to insect, *ρ* is the relative risk of transmission from a subclinical compared to a clinical bovine, *γ* is the virus inactivation rate, *p*_*C*_ is the proportion of cattle that develop clinical disease and 1/*r*_*C*_ and 1/*r*_*S*_ are the mean durations of infectiousness for clinical and subclinical animals, respectively, all of which were estimated in the present study, and *a, m* and *µ* are the biting rate, vector to host ratio and vector mortality rate, respectively. The formal derivation of this expression, (4), is given in S1 Text.

Replicated Latin hypercube sampling was used to compute the median and 95% prediction interval for *R*_0_ for each insect species [45]. Parameters were sampled either from their marginal posterior distributions derived in the present study (*b, β, ρ, γ, p*_*C*,_ 1/*r*_*C*_ and 1/*r*_*S*_; see Tables 1 and S2) or uniformly from plausible ranges (*a, m* and *μ*; see Gubbins [45], their Table 1). The mean duration of infection for clinical animals (1/*r*_*C*_) is based on detection of virus or viral DNA in skin, while that for subclinical animals (1/*r*_*S*_) is based on detection of viral DNA in blood (S2 Table).

## Data availability

The authors declare that the main data supporting the findings of this study are available within the article and its Supplementary Information files.

## Code availability

The code and the data used are available online for readers to access with no restriction at

https://github.com/SimonGubbins/LSDVAcquisitionAndRetentionByInsects.

## Funding information

This study was funded by the Biotechnology and Biological Sciences Research Council (BBSRC) [grant code: BB/R002606/1] and by MSD Animal Health. SB was supported by the Wellcome Trust (200171/Z/15/Z to LA); KED, SG, LA and PMB were supported by BBSRC strategic funding (BBS/E/I/00007033, BBE/I/00007036, BBS/E/I/00007037, BBS/E/I/00007038 and BBS/E/I/00007039).

## Conflict of interest

None to report.

## Acknowledgements

The authors are grateful to the Animal Services and the Non-Vesicular Reference Laboratory staff at The Pirbright Institute for their invaluable assistance in the running of the animal studies and sample assays, and to Dr. Lara Harrup for useful technical advice.

## Author contribution

P.M.B., S.G., K.E.D., P.C.H., J.A. and A.J.W. conceptualised the study; B.S.B., L.A., S.B., C.S. and C.B. contributed to the design of the experiments. S.B., W.L., A.V.D., Z.L., E.D., J.S. and M.W. prepared the insects; B.S.B., P.M.B., I.R.H., N.W. and K.E.D. carried out the cattle experiments including collection and preparation of samples. B.S.B. performed the laboratory assays and data acquisition. S.G. performed the statistical analysis and mathematical modelling. B.S.B., S.G. and P.M.B. drafted the paper. All authors discussed the results and commented on the manuscript.

## Supporting information

**Data S1**. Clinical observations and levels of viral DNA and infectious virus in samples taken from calves infected with lumpy skin disease virus and the outcome when blood-feeding insects were allowed to feed on them.

**Fig S1**. Clinical characterisation of experimental challenge of cattle with lumpy skin disease virus.

**S2 Fig**. A higher proportion of insects are positive for lumpy skin disease viral DNA after feeding on a clinical compared to a subclinical animal.

**S3 Fig**. The proportion of blood-feeding insects positive for lumpy skin disease viral DNA differs over time post challenge.

**S4 Fig**. The median amount of lumpy skin disease viral DNA in homogenised whole insects was the same over time post feeding in three out of four species tested.

**S1 Table**. Deviance information criterion (DIC) for models assessing variation amongst insect species in the virus inactivation rate, the probability of transmission of lumpy skin disease virus from bovine to insect and the relative risk of transmission from a subclinical bovine.

**S2 Table**. Parameters for the duration of latent and infectious periods for lumpy skin disease virus in cattle.

**S1 Text**. Modelling the transmission of lumpy skin disease virus by haematophagus insects.

## Notes

https://github.com/SimonGubbins/LSDVAcquisitionAndRetentionByInsects

### Competing Interest Statement

The authors have declared no competing interest.

