## Supplementary material for "Quantifying and modelling the acquisition and retention of lumpy skin disease virus by haematophagus insects reveals clinically but not subclinically-affected cattle are promoters of viral transmission and key targets for control of disease outbreaks": Figure S1 (from another version of this preprint: version 2, 2020-08-26)

A) Clinical signs of lumpy skin disease

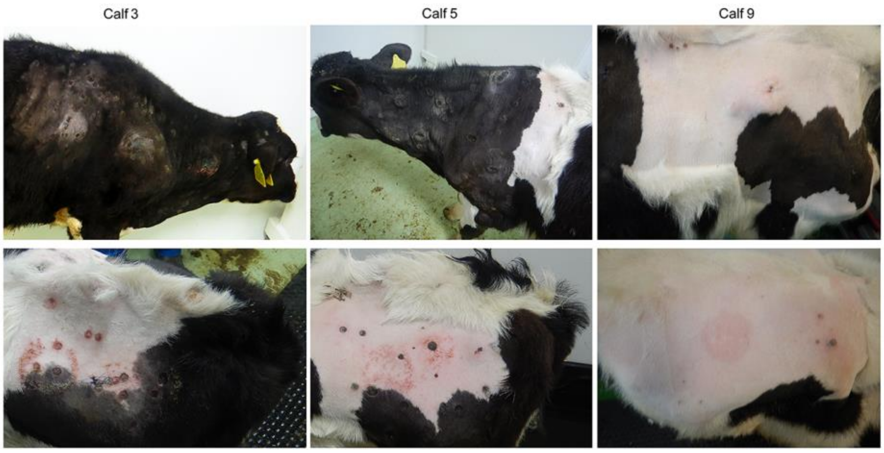


B) Cutaneous nodules at the injection site

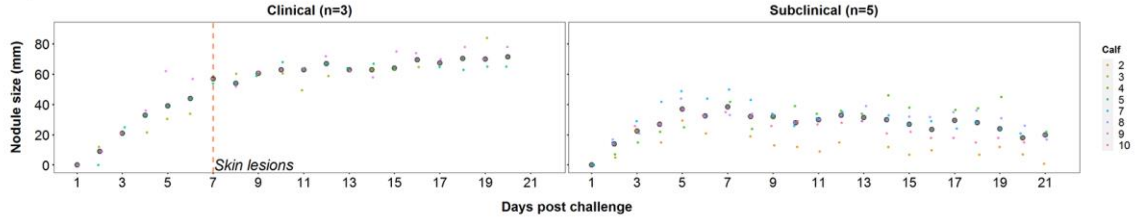


C) Rectal temperature

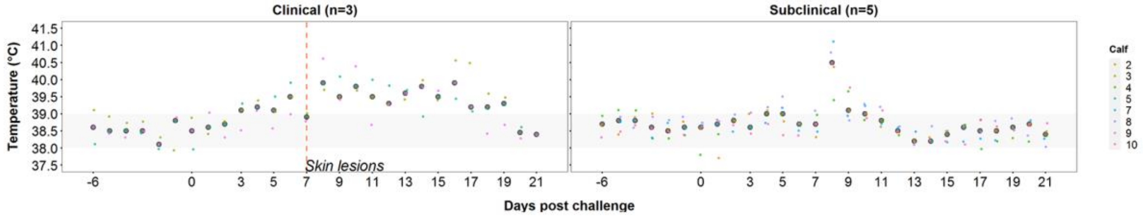


D) Cumulative score for lymphadenopathy

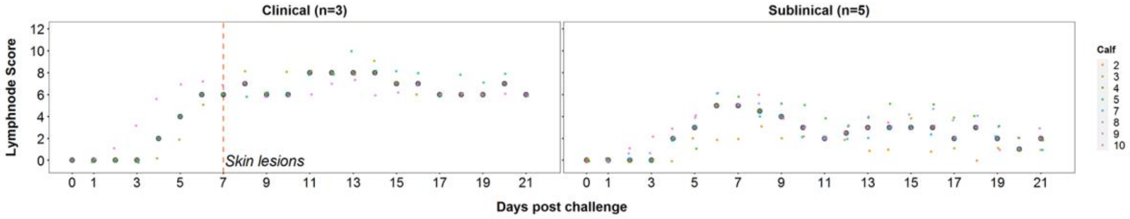


E) Serum antibody levels

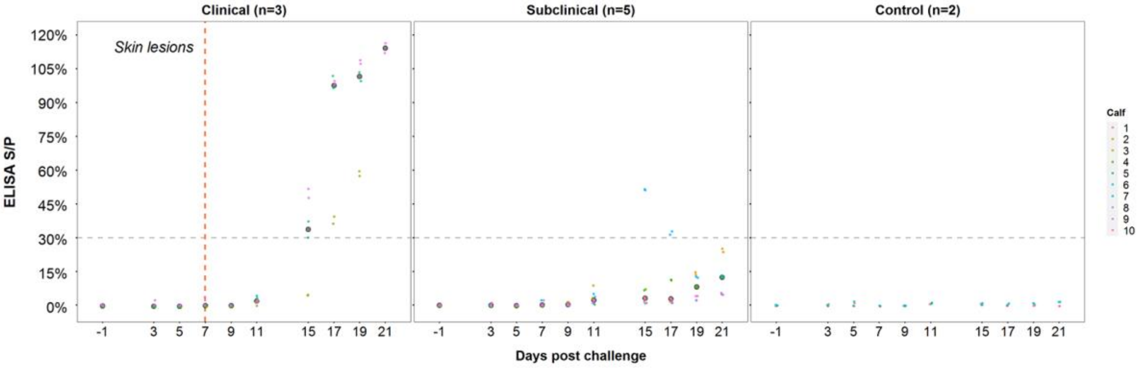

**Figure S1. Clinical characterisation of experimental challenge of cattle with lumpy skin disease virus.** A) LSDV causes mild to severe multifocal necrotising dermatitis in a subset of inoculated calves. Top row: Photographs of the right shoulder (calves 3 and 9) or left shoulder (calf 5) at 19 dpc (calf 3), 18 dpc (calf 5) or 16 dpc (calf 9). Calves 3 and 5 have numerous large (up to 5cm diameter) necrotic lesions visible. In comparison calf 9 has fewer and smaller lesions - three cutaneous lesions (dorsal) and one subcutaneous lesion (central), in addition to a biopsy site with blue suture material. Bottom row: Photographs of the left rump of calves 3, 5 and 9 at 19 dpc (calf 3), 18 dpc (calf 5) and 15 dpc (calf 9). Calves 3 and 5 have approximately 10 sunken, necrotic, well circumscribed cutaneous lesions up to 1.5cm in diameter. In comparison calf 9 has three lesions. In all three calves circular red areas approximately 6.4 cm in diameter can be observed on the skin. These correspond to areas where Culicoides nubeculosus fed on the calves. B) Diameter of the nodules formed at the challenge site after the intradermal inoculation of LSDV. C) Daily rectal temperature of calves challenged with LSDV. The shaded area is the range of normal temperature values for cattle (1). D) Cumulative score of superficial lymph nodes. The superficial cervical and the iliac lymph nodes were scored daily to make a cumulative score [Scores: 0=normal, 1=enlarged on palpation, 2=very enlarged on palpation, 3=enlarged on visual inspection]. Each graph shows the values for individual animals (coloured circles) and the median (grey circles). The orange line marks the start of generalised secondary skin lesion in animals with clinical disease. E) LSDV antibodies are detected in the serum of challenged calves. Each plot shows the individual sample/positive (S/P) ratio (%) over time with the median value (grey circle; not for control animals) and for each calf in duplicate (coloured circles). The orange line marks the time generalised secondary skin lesions first appeared in animals with clinical disease. The dotted line indicates the threshold for positivity (30%) recommended by the manufacturer.
