## Supplementary material for "Quantifying and modelling the acquisition and retention of lumpy skin disease virus by haematophagus insects reveals clinically but not subclinically-affected cattle are promoters of viral transmission and key targets for control of disease outbreaks": Table S1 (from another version of this preprint: version 2, 2020-08-26)

**Table S1.** Deviance information criterion (DIC) for models assessing variation amongst insect species in the virus inactivation rate, the probability of transmission of lumpy skin disease virus from bovine to insect and the relative risk of transmission from a subclinical bovine.

| dose-independent model* | | | DIC† | dose-response model* | | | DIC† | |
|---|---|---|---|---|---|---|---|---|
| inactivation rate | probability of transmission | relative risk of transmission | | inactivation rate | intercept | slope | blood‡ | skin‡ |
| common | common | no difference | 1335.3 | common | common | zero | 1335.3 | 1335.3 |
| varies | common | no difference | 1334.6 | common | common | common | 874.2 | 684.6 |
| common | varies | no difference | 1338.6 | common | varies | common | 871.1 | 682.0 |
| varies | varies | no difference | 1340.2 | common | common | varies | 869.0 | 681.4 |
| common | common | common | 1025.5 | common | varies | varies | 868.9 | 680.7 |
| varies | common | common | **1021.0** | varies | common | zero | 1334.6 | 1334.6 |
| common | varies | common | 1026.8 | varies | common | common | **863.8** | **675.1** |
| varies | varies | common | 1025.9 | varies | varies | common | 864.0 | 676.5 |
| common | common | varies | 1025.3 | varies | common | varies | 863.9 | 677.1 |
| varies | common | varies | 1019.5¶ | varies | varies | varies | 863.9 | 677.4 |
| common | varies | varies | 1025.5 | | | | | |
| varies | varies | varies | 1024.5 | | | | | |

* “common”: parameter common to all insect species; “varies”: parameter varies amongst insect species; “no difference”: risk of transmission does not differ between clinical and subclinical animals; “zero”: parameter fixed at zero

† a model with a lower DIC is preferred to one with higher DIC

‡ level of viral DNA in blood or skin used as the proxy measure for infectiousness

¶ although this model has a smaller DIC than the one shown in bold, the difference is less than two and the simpler model was preferred as it has the smaller number of parameters
