## Supplementary material for "Quantifying and modelling the acquisition and retention of lumpy skin disease virus by haematophagus insects reveals clinically but not subclinically-affected cattle are promoters of viral transmission and key targets for control of disease outbreaks": Table S2 (from another version of this preprint: version 2, 2020-08-26)

**Table S2.** Parameters for the duration of latent and infectious periods for lumpy skin disease virus in cattle.

| proxy measure of infectiousness | shape parameter ($k_i$)* | mean ($1/r_i$; days)* |
|---|---|---|
| *latent period (days)* | | |
| skin lesions† | 38.0 (10.9, 103.3) | 7.6 (6.6, 8.9) |
| viraemia (PCR)† | 3.8 (2.0, 6.9) | 5.4 (4.2, 7.1) |
| viraemia (PCR)‡ | 3.9 (1.4, 9.1) | 5.5 (3.9, 8.3) |
| *infectious period (days)* | | |
| skin biopsies† | 11.4 (2.8, 38.2) | 23.6 (17.1, 38.0) |
| viraemia (PCR)† | 3.4 (1.3, 7.6) | 19.0 (13.7, 30.4) |
| viraemia (PCR)‡ | 2.5 (0.9, 5.7) | 15.2 (10.2, 27.2) |

* posterior median (95% credible interval)

† clinically-affected animals only

‡ subclinical animals only
