## Supplementary material for "Quantifying and modelling the acquisition and retention of lumpy skin disease virus by haematophagus insects reveals clinically but not subclinically-affected cattle are promoters of viral transmission and key targets for control of disease outbreaks": Text S1 (from another version of this preprint: version 2, 2020-08-26)

**Text S1. Modelling the transmission of lumpy skin disease virus by haematophagus insects**

### S1.1 Probability of transmission from bovine to insect and virus inactivation rate

For each insect, the clinical status of the donor calf (i.e. clinical or subclinical), the number of days post feeding at which the insect was tested and whether the insect was positive for lumpy skin disease virus (LSDV) DNA (defined as any viral DNA detected) were used to estimate the probability of transmission from bovine to insect and the virus inactivation rate.

The likelihood for the data is given by,

$$L = \prod_i p_i^{\delta_i} (1 - p_i)^{1-\delta_i}, \tag{S.1}$$

where $\delta_i$ is an indicator for whether insect $i$ was negative ($\delta_i$=0) or positive ($\delta_i$=1) for viral DNA when tested for LSDV and $p_i$ is the probability that insect $i$ was positive when tested. If insect $i$ fed on a clinically affected animal. this is given by,

$$p_i = \beta_{s_i} \exp(-\gamma_{s_i} t_i), \tag{S.2}$$

while if it fed on a subclinical donor, it is given by,

$$p_i = \rho_{s_i} \beta_{s_i} \exp(-\gamma_{s_i} t_i), \tag{S.3}$$

Equations (S.2) and (S.3) combine: (i) the species-specific probability that an insect became infected ($\beta_s$ or $\rho_s\beta_s$, depending on the status of the donor calf, where $\beta_s$ is the probability of transmission from a clinical bovine to an insect and $\rho_s$ is the relative risk of transmission from a subclinical donor compared to a clinical one); and (ii) the probability that the insect was still positive when it was tested at $t_i$ days post feeding ($\exp(-\gamma_s t_i)$, where $\gamma_s$ is the species-specific virus inactivation rate).

Differences amongst insect species in the virus inactivation rate, probability of transmission from bovine to insect and relative risk of transmission were incorporated in the model by assuming hierarchical structure in the parameters, so that,

$$\begin{aligned} \gamma_s &\sim \text{Gamma}(k_\gamma, \mu_\gamma), \\ \beta_s &\sim \text{Beta}(a_\beta, b_\beta), \\ \rho_s &\sim \text{Gamma}(k_\rho, \mu_\rho), \end{aligned} \quad \text{(S.4)}$$

where $k_\gamma$ and $k_\rho$ are shape parameters and $\mu_\gamma$ and $\mu_\rho$ are means for a gamma distribution and $a_\beta$ and $b_\beta$ are the parameters for a beta distribution. If and how the parameters varied amongst species was explored by considering the fit of the model when no, one, two or three of the parameters varied amongst the insect species. In total, twelve models were considered (see Table S1).

Hierarchical priors were used for those parameters that differed amongst species and exponential priors (with mean 1) were used for the higher-order parameters in the hierarchical distributions. Where the parameters were common to all species, an exponential prior (with mean 100) was used for $\gamma$ or $\rho$ and a uniform prior (with range (0,1)) was used for $\beta$.

Samples from the joint posterior density were generated using an adaptive Metropolis scheme (Haario et al. 2001), modified so that the scaling factor was tuned during burn-in to ensure an acceptance rate of between 20% and 40% for more efficient sampling of the target distribution (Andrieu & Thoms 2008). Two chains of 5,000,000 iterations were run, with the preceding 1,000,000 iterations discarded to allow for burn-in of the chain. The chains were then thinned (taking every 500th sample) to reduce autocorrelation amongst the samples. The adaptive Metropolis scheme was implemented in Matlab (version R2019b; The Mathworks Inc.) and the code is available online at https://github.com/SimonGubbins/LSDVAcquisitionAndRetentionByInsects. Convergence of the scheme was assessed visually and by examining the Gelman-Rubin statistic provided in the coda package (Plummer et al. 2006) in R (version 3.6.1; R Core Team 2019). Different models for the variation amongst species in virus inactivation and probability of transmission from bovine to insect were compared using the deviance information criterion (DIC) (Spiegelhalter et al. 2002).

### S1.2 Dose-response relationship

For each insect, the level of lumpy skin disease viral DNA in blood ($\log_{10}$ copies/ml) or skin ($\log_{10}$ copies/mg) for the calf at the time when the insect fed, the number of days post feeding at which the insect was tested and whether or not the insect was positive for LSDV DNA (defined as any viral DNA detected) were used to investigate the dose-response relationship for the probability of transmission. Only data for insects which fed on the clinically affected donors were included in this analysis.

The likelihood for the data is given by equation (S.1), but the probability that insect $i$ was positive is given by,

$$p_i = \beta_i \exp(-\gamma_{s_i} t_i), \tag{S.5}$$

where $\beta_i$ is the probability of transmission from bovine to insect for insect $i$, $\gamma_s$ is the virus inactivation rate for species $s$ and $t_i$ is the number of days post feeding at which insect $i$ was tested. The dose-response relationship between the probability of transmission from bovine to insect (i.e. $\beta_i$) and level of viral DNA was described by,

$$\log\left(\frac{\beta_i}{1-\beta_i}\right) = a_{s_i} + b_{s_i} V_{c_i}(t_f^{(i)}), \tag{S.6}$$

where $a_s$ and $b_s$ are (insect) species-specific dose-response parameters and $V_j(t)$ is the level of viral DNA in calf $j$ at $t$ days post infection, $c_i$ is the calf on which insect $i$ fed and $t_f^{(i)}$ is the time of feeding for insect $i$.

Variation amongst insect species in virus inactivation rates ($\gamma$) and dose response ($a$, $b$) was incorporated in the model by assuming hierarchical structure in the parameters, so that,

$$\begin{aligned} \gamma_s &\sim \text{Gamma}(s_\gamma, \mu_\gamma), \\ a_s &\sim \text{Normal}(\mu_a, \sigma_a^2), \\ b_s &\sim \text{Normal}(\mu_b, \sigma_b^2), \end{aligned} \quad \text{(S.7)}$$

where $\mu_\gamma$, $\mu_a$ and $\mu_b$ are the distribution means, $s_\gamma$ is the shape parameter for the gamma distribution and $\sigma_a$ and $\sigma_b$ are the standard deviations for the normal distributions. Five possibilities were considered for the parameters relating titre to the probability of transmission: (i) independent of titre (i.e. $a$ common to all species and $b$=0); (ii) $a$ and $b$ were common to all species; (iii) $a$ varied amongst species and $b$ was common to all species; (iv) $a$ was common to all species and $b$ varied amongst species; and (v) $a$ and $b$ varied amongst species. In addition, two possibilities were considered for the the virus inactivation rate: (i) common to all species; or (ii) varies amongst species. Consequently, ten models were considered (Table S1).

Where the parameters were common to all species an exponential prior (with mean 100) was used for $\gamma$ and normal priors (with mean 0 and standard deviation 10) were used for $a$ or $b$. Hierarchical priors were used for those parameters that differed amongst species. For the higher-order parameters in the hierarchical distributions, an exponential prior (with mean 1) was used for $\mu_\gamma$ and normal priors (with mean 0 and standard deviation 10) were used for $\mu_a$ and $\mu_b$, while exponential priors with mean 1 were used for $s_\gamma$, $\sigma_a$ and $\sigma_b$.

Samples from the joint posterior density were generated using an adaptive Metropolis scheme as described in section S1.1 above. Two chains of 5,000,000 iterations were run, with the preceding 1,000,000 iterations discarded to allow for burn-in of the chain. The chains were then thinned (taking every 500th sample) to reduce autocorrelation amongst the samples.

The different models for the relationship between titre and infection were compared using the deviance information criterion (Spiegelhalter et al. 2002). The two proxy measures for infectiousness (i.e. level of viral DNA in blood or skin) were compared by computing posterior predictive *P*-values for each insect. Specifically, the joint posterior distribution for the model was sampled and the probability that each insect was positive for viral DNA computed. Whether or not the insect was positive when tested was then simulated and the observed outcomes were compared to the simulated ones. This was repeated multiple times and the proportion of samples for which the observed and simulated outcomes matched was computed (i.e. the posterior predictive *P*-value).

#### S1.3 Probability of transmission from insect to bovine

Previous studies have considered the transmission of LSDV from insect to bovine for the four insect species used in the present study (Chihota et al. 2001, 2003; Sohier et al. 2019). The results of these earlier studies were analysed in light of the results from the present study to estimate the probability of transmission from insect to bovine for each species.

***Aedes aegypti & Culex quinquefasciatus.*** A batch of insects was fed on a clinically affected bovine (*Ae. aegypti*) or a blood-virus mix via a membrane (*Cx. quinquefasciatus*) and, subsequently, allowed to refeed on a naïve bovine, with each recipient fed on only once (Chihota et al. 2001, 2003). The likelihood for the data for each species is

$$L = \prod_j q_j^{I_j} (1 - q_j)^{1 - I_j}, \tag{S.8}$$

where $I_j$ is a variable indicating whether ($I_j$=1) or not ($I_j$=0) transmission occurred when insects were allowed to refeed on naïve animal $j$ at time $t_j$ post initial feed and

$$q_j = 1 - (1 - b\beta \exp(-\gamma t_j))^{n_j}, \tag{S.9}$$

is the probability of transmission for the challenge. The probability, (S.9), is the probability that at least one insect (out of the $n_j$ refeeding) transmitted LSDV, where the probability that an individual insect will transmit is the product of the probabilities that it became infected during the initial feed ($\beta$), was still infected at refeeding ($\exp(-\gamma t_j)$) and that it subsequently transmitted LSDV at refeeding ($b$). The marginal posterior densities for each species obtained in section S4.1 were used as informative priors for $\beta$ and $\gamma$, while a non-informative prior (uniform with range (0,1)) was used for $b$.

Samples from the joint posterior density were generated using an adaptive Metropolis scheme similar to that described in section S1.1 above. Two chains of 50,000 iterations were run, with the preceding 25,000 iterations discarded to allow for burn-in of the chain. The chains were then thinned (taking every 5th sample) to reduce autocorrelation amongst the samples.

***Culicoides nubeculosus.*** The experimental design for *C. nubeculosus* (Chihota et al. 2003) was the same as described above for *Ae. aegypti*. However, the number of insects refeeding was not reported, yet this is required when estimating the probability of transmission from insect to bovine (see equation (S.9)). Accordingly, the numbers of insects refeeding in each batch were included in a data augmentation step in the Bayesian framework (i.e. as nuisance parameters). In this case, the number of insects refeeding was drawn from a binomial distribution for this species, so that

$$n_j \sim \text{Binomial}(N_j, 1-\exp(-at_j)), \qquad \text{(S.10)}$$

where $N_j$ is the number of insects used for challenge $j$ (assumed to be 100, which is approximately the maximum number of insects fed), $a$ is the biting rate and $t_j$ is the time between feeding on the donor and refeeding on the recipient. A uniform prior with range (0,0.4)) was used for $a$ (based on the duration of the gonotrophic cycle of *Culicoides* biting midges; Birley & Boorman 1982; Braverman et al. 1985; Mullens et al. 2004).

Samples from the joint posterior density were generated using an adaptive Metropolis scheme similar to that described in section S1.1 above. Two chains of 500,000 iterations were run, with the preceding 100,000 iterations discarded to allow for burn-in of the chain. The chains were then thinned (taking every 50th sample) to reduce autocorrelation amongst the samples.

***Stomoxys calcitrans.*** Two studies have examined transmission of LSDV by *S. calcitrans* (Chihota et al. 2003; Sohier et al. 2019). The same experimental design described above for *C. nubeculosus* was used by Chihota et al. (2003), so the same likelihood applies for these data (i.e. including the data augmentation step). In the experiments conducted by Sohier et al. (2019), batches of insects were allowed to feed on an infected donor for ten minutes and, one hour later, were allowed to refeed on a naïve recipient for ten minutes. For each batch this was repeated one, two or three times at daily intervals.

The probability that an animal became infected (with onset of viraemia on day $T$ post initial feed) is given by,

$$p_{POS} = 1 - \prod_j (1 - p_j), \qquad \text{(S.11)}$$

where

$$p_j = \sum_k \left\{ q_{jk} \prod_{i=1}^{k-1} (1 - q_{ji}) \right\} \int_{T-t_{jk}-1}^{T-t_{jk}} f(\tau)\, \mathrm{d}\tau, \tag{S.12}$$

is the probability of transmission for batch of insects $j$. Here $q_{jk}$ is the probability of transmission during the $k$th feed of batch $j$ (on day $t_{jk}$) and $f$ is the probability density function for the gamma-distributed latent period for LSDV (with mean $1/r_E$ and shape parameter $k_E$). The first term in the summation (in braces) is the probability the animal became infected on the $k$th feed (and was not infected in any of the previous feeds). The second term (the integral) is the probability that, if the animal became infected on the $k$th feed, the onset of viraemia would occur on day $T$ after the initial feed.

The probability of transmission during feed $k$ for batch of insects $j$ is given by

$$q_{jk} = 1 - \left( 1 - b \sum_{i=1}^{k} \beta \exp(-\gamma(t_{ji} - t_{j1} + \delta)) \right)^{n_{jk}}. \tag{S.13}$$

This is the probability that at least one insect (out of the $n_{jk}$ refeeding) transmitted LSDV. The probability that an individual insect will transmit is the probability that it became infected during the any of preceding feeds on the infected donor and was still infected at refeeding ($\beta\exp(-\gamma(t_{ji}-t_{j1}+\delta))$[1], where $\delta$ is the time interval between feeding on the donor and refeeding on the recipient), multiplied by the probabilty that it transmitted LSDV at refeeding ($b$). Because the number of insects refeeding was not known, these were included as nuisance parameters (cf. *C. nubeculosus* above), with

$$n_{jk} \sim \mathrm{Binomial}(N_j, 1 - \exp(-a\delta)), \tag{S.14}$$

where $N_j$ is the number of insects in batch $j$ and $a$ is the biting rate.

[1] If the donor animal was subclinically-affected, the $\beta$ in this expression is replaced by $\rho\beta$ (see section S2.1).

The probability that an animal remained uninfected is given by the probability that transmission did not occur at any of the feeds for any of the batches of insects, that is

$$p_{NEG} = \prod_j \prod_k (1 - q_{jk}), \tag{S.15}$$

where $q_{jk}$ is as defined in equation (S.13).

The likelihood for the Sohier data is

$$L = \prod_a p_{NEG}^{1-I_a} p_{POS}^{I_a}, \tag{S.16}$$

where $\delta_a$ is a variable indicating whether ($I_a$=1) or not ($I_a$=0) transmission occured to animal $a$ and $p_{POS}$ and $p_{NEG}$ are the probabilities that an animal became infected or remained uninfected (given by equations (S.12) and (S.15)), respectively. The overall likelihood for the *S. calcitrans* data is the product of that for the Chihota data, (S.8)-(S.10), and that for the Sohier data, (S.16).

The marginal posterior densities for $\beta$, $\rho$ and $\gamma$ derived for *S. calcitrans* in section S4.2 were used as informative priors for these parameters. A non-informative prior (uniform with range (0,1)) was used for $b$. A gamma prior (with mean 2.26 and shape parameter 2.27) was used for the biting rate $a$, based on an interval between blood meals of 4 to 72 hours (Salem et al. 2012; Baldacchino et al. 2013). The marginal posterior densities obtained from analysis of previous challenge experiments and the present study (see Text S5 for details) were used as informative priors for the mean and shape parameter for the latent period distribution (Table S5).

Samples from the joint posterior density were generated using an adaptive Metropolis scheme similar to that described in section S1.1 above. Two chains of 10,000,000 iterations were run, with the preceding 1,000,000 iterations discarded to allow for burn-in of the chain. The chains were then thinned (taking every 1000th sample) to reduce autocorrelation amongst the samples.

### S1.4 Latent and infectious periods for lumpy skin disease virus in cattle

The mean ($1/r_i$) and shape parameter ($k_i$) for the gamma-distributed latent ($i=E$) and infectious ($i=I$) periods for LSDV in clinically-affected cattle have been estimated previously (Gubbins 2019) using data from experimental infections of cattle (Chihota et al. 2001; Tuppurainen et al. 2005; Babiuk et al. 2008). These estimates were updated (Table S2) using the outcome of the present study for two proxy measures of infectiousness: detection of viral DNA in blood; and detection of virus or viral DNA in skin biopsies. This was done using the methods described in Gubbins (2019), but with the marginal posterior densities from the earlier analysis providing informative priors for the mean and shape parameters.

To estimate the corresponding parameters for subclinical cattle (Table S5), detection of viral DNA in blood in experimentally-infected cattle as reported in published data (Tuppurainen et al. 2005; Möller et al. 2019; Sohier et al. 2019) and the present study was used as a proxy measure of infectiousness (as the animals did not have skin lesions). In this case, the methods described in Gubbins (2019) were used, including non-informative priors.

### S1.5 Proportion of cattle developing generalised disease

The proportion of cattle developing generalised disease ($p_C$) was estimated based on the numbers of cattle that developed generalised disease following challenge in published experiments: 5 out of 14, 4 out of 25 and 8 out of 11 (Carn & Kitching 1995); 3 out of 6

(Möller et al. 2019); 2 out of 6 (Tuppurainen et al. 2005); 6 out of 7 (Babiuk et al. 2008); 1 out of 4, 3 out of 5 and 4 out of 5 (Sohier et al. 2019); and 3 out of 8 (present study).

Parameters were estimated in a Bayesian framework. The likelihood for the data is

$$L = \prod_i \binom{N_i}{C_i} p_i^{C_i} (1 - p_i)^{N_i - C_i},$$

where $C_i$ and $N_i$ are the number of cattle developing generalised disease and the number of challenged cattle, respectively, and $p_i$ is the expected proportion of cattle developing generalised disease in experiment $i$. To allow the $p_i$s to vary amongst experiments, a hierarchical structure was assumed, such that,

$$p_i \sim \mathrm{Beta}(\alpha_p, \beta_p),$$

where $\alpha_p$ and $\beta_p$ are hierarchical parameters. Exponential priors (with mean 100) were assumed for $\alpha_p$ and $\beta_p$. Samples from the joint posterior distribution were generated by Markov chain-Monte Carlo methods implemented in OpenBUGS (version 3.2.3). Two chains each of 100,000 samples were run, with the preceding 20,000 iterations discarded to allow for burn-in. The chains were then thinned (selecting every 20th sample) to reduce autocorrelation. The posterior median (95% credible interval (CI)) for $\alpha_p$ and $\beta_p$ were 38.5 (3.4, 196.4) and 49.0 (3.6, 256.7), respectively, with a corresponding posterior median for $p_C$ of 0.44 (95% CI: 0.34-0.55).

A simpler model in which the proportion of cattle developing generalised disease was common to all experiments was also considered. However, this yielded a poorer fit to the data, as judged by the deviance information criterion: 44.0 ($p$ varying) vs 48.0 ($p$ common).

### S1.6 Basic reproduction number for lumpy skin disease virus in cattle

The basic reproduction number, $R_0$, is calculated as the dominant eigenvalue, $r(K)$, of the next-generation operator, $K$ (Diekmann & Heesterbeek, 2000; Heffernan et al. 2005). For lumpy skin disease virus, the next generation operator is a matrix, **K**, whose elements ($K_{ij}$) are the expected number of infections of type $i$ (either clinical cattle ($C$), subclinical cattle ($S$) or vectors ($V$)) arising from a single infected individual of type $j$ (cf. Gubbins 2019). In this case, **K** can be written as,

$$\mathbf{K} = \begin{pmatrix} K_{CC} & K_{CS} & K_{VC} \\ K_{SC} & K_{SS} & K_{VS} \\ K_{VC} & K_{SV} & K_{VV} \end{pmatrix}. \tag{S.17}$$

Some elements of **K** are straightforward to derive: there is assumed to be no direct transmission between hosts or vectors (i.e. $K_{CC}$=$K_{CS}$=$K_{SC}$=$K_{SS}$=$K_{VV}$=0). The remaining elements, representing transmission from vector to host ($K_{CV}$, $K_{SV}$) or host to vector ($K_{VC}$, $K_{VS}$), are computed as follows.

*Transmission from vector (V) to host (C or S):* After an insect feeds on an infected bovine, it remains infected (and infectious) until the virus becomes inactivated or the insect dies, a period which lasts on average $1/(\gamma+\mu)$ days, where $\gamma$ is the virus inactivation rate and $\mu$ is the vector mortality rate. During this time the infected insect will bite susceptible cattle $a$ times per day and a proportion, $b$, of these bites will result in a newly infected animal. Of these, a proportion, $p_C$ will go on to develop generalised disease, while the remainder, $p_S$=1-$p_C$, will remain subclinical. Hence, the expected number of infected cattle of each type per infected insect ($K_{iV}$, $i$=$C$,$S$) is given by,

$$K_{iV} = \frac{ba}{\gamma+\mu} p_i. \tag{S.18}$$

*Transmission from host (C or S) to vector (V)):* An infected bovine remains infectious for $1/r_i$ days. During this time the it will be bitten by susceptible insects on average $m \times a$ times per day, a proportion, $\beta_i$ (where $\beta_C = \beta$ and $\beta_S = \rho\beta$, where $\beta$ is the probability of transmission from bovine to insect for a clinically-affected animals and $\rho$ is the relative risk of transmission from a subclinical compared to a clinical bovine), of which will result in a newly infected vector. Consequently, the expected number of infected insects per infected bovine ($K_{Vi}$) is given by,

$$K_{Vi} = \frac{\beta_i m a}{r_i}. \tag{S.19}$$

Some linear algebra shows that the dominant eigenvalue of **K** (i.e. $R_0$) is,

$$R_0 = \sqrt{K_{CV} K_{VC} + K_{SV} K_{VS}}, \tag{S.20}$$

which, on substituting the expressions in (S.18) and (S.19) becomes,

$$R_0 = \sqrt{\frac{b\beta m a^2}{(\mu + \gamma)} \left( p_C \frac{1}{r_C} + (1 - p_C)\rho \frac{1}{r_S} \right)}. \tag{S.21}$$
